# Sparse Graphical Models for Functional Connectivity Networks: Best Methods and the Autocorrelation Issue

**DOI:** 10.1101/128488

**Authors:** Yunan Zhu, Ivor Cribben

## Abstract

Sparse graphical models are frequently used to explore both static and dynamic functional brain networks from neuroimaging data. However, the practical performance of the models has not been studied in detail for brain networks. In this work, we have two objectives. First, we compare several sparse graphical model estimation procedures and several selection criteria under various experimental settings, such as different dimensions, sample sizes, types of data, and sparsity levels of the true model structures. We discuss in detail the superiority and deficiency of each combination. Second, in the same simulation study, we show the impact of autocorrelation and whitening on the estimation of functional brain networks. We apply the methods to a resting-state functional magnetic resonance imaging (fMRI) data set. Our results show that the best sparse graphical model, in terms of detection of true connections and having few false-positive connections, is the smoothly clipped absolute deviation (SCAD) estimating method in combination with the Bayesian information criterion (BIC) and cross-validation (CV) selection method. In addition, the presence of autocorrelation in the data adversely affects the estimation of networks but can be helped by using the CV selection method. These results question the validity of a number of fMRI studies where inferior graphical model techniques have been used to estimate brain networks.

## Introduction

**T**o thoroughly understand brain function, researchers have begun to map the functional network. In this study, the emphasis is on studying the interaction of brain regions, as a great deal of neural processing is performed by an integrated network of several brain regions. This is sometimes referred to as functional integration. In the analysis of functional magnetic resonance imaging (fMRI), positron emission tomography (PET), electroencephalography (EEG), and magnetoencephalography (MEG) time series data, functional connectivity (FC) is the name given to the interaction, correlation, or dependence between signals observed from spatially remote brain regions (Friston et al., 1993). FC is a measure of dependence or “relatedness” but does not comment on how the dependence is mediated. It is sometimes referred to as undirected association. Estimating the FC between predefined brain regions or voxels allows for the characterization of interregional neural interactions during particular experimental tasks or merely from spontaneous brain activity while subjects are being scanned at rest. Using fMRI, researchers have been able to create maps of FC with distinct spatial distributions of temporally correlated brain regions called functional networks.

The accurate estimation of FC networks is important as it has been shown that neurological disorders, such as schizophrenia, depression, anxiety, dementia, and autism, disrupt the FC or structural properties of the brain (Menon, 2011). However, it is still unclear whether the disruptions are the cause or consequence of the disorder. The estimation of FC and linking its structure to disorders is a good starting point for treatment of the disorder. Calhoun et al. (2009) investigated the link between FC and schizophrenia with the objective of finding biomarkers for the disorder. Buckner et al. (2009) found different static FC network structures in subjects with Alzheimer’s disease compared with healthy subjects.

In general, to estimate FC, we carry out two major steps: we calculate the average voxel time series from prespecified brain regions and estimate the dependence between the time series. Typically, higher similarities of the time courses between any given pair of brain regions indicate a higher chance of an FC between those nodes. The simplest methods for estimating FC include the sample correlation matrix (CM) and the sample partial CM (PCM). However, these methods are deficient since they are rarely estimated to be exactly zero even if the data are independent. Alternatively, the FC or functional network can also be represented by an undirected graphical model. In this study, the nodes of the undirected graph represent the functional regions of the brain, and the edges of the graph represent the connections between those functional nodes. The estimate of a precision matrix (or inverse covariance matrix) can be illustrated using an undirected graph. A nonzero estimated entry of the precision matrix corresponds to an edge of the undirected graph, while an absence of an edge between two functional nodes indicates conditional independence between them. Also, the thickness of the edge of the undirected graph indicates the strength of the conditional connection between the corresponding functional nodes. Hence, the elements of a standardized precision matrix are equivalent to partial correlations. A sparse undirected graph, where some edges are set exactly to zero, is usually preferred for its simplicity and ease of interpretation. Smith et al. (2011) pointed out that correlation does not necessarily imply either the causality of a connection or whether it is direct. However, the partial correlation can correctly estimate the true network, which captures direct connections only. Smith et al. (2011) conclude that with respect to estimating FC networks, partial correlations are within the “Top 3” methods. Moreover, the partial correlation and the regularized precision matrix are very sensitive in detecting the network connections on good-quality fMRI data.

Several methods for estimating FC or brain networks using sparse graphical models have been proposed. The estimation methods are all based on penalized log-likelihood methods, which apply a regularization parameter to control the sparsity of the graph. For example, the graphical lasso or glasso (Friedman et al., 2007b) is a method known for its computational speed, ease of implementation, and its production of a sparse undirected graph. A newly developed algorithm (Mazumder and Hastie, 2012), called the DP-glasso, differs from glasso in that it solves for the precision matrix O and not the covariance matrix estimated by glasso. The bootstrap graphical lasso (BG) (Cribben et al., 2013) estimating method, inspired by the stability selection (SS) technique of Meinshausen and Bühlmann (2010), combines both glasso and the bootstrap resampling procedure (Efron and Tibshirani, 1994) to improve the estimation performance. The estimating methods, the adaptive lasso (AL), and the smoothly clipped absolute deviation (SCAD) are modifications of glasso in the sense that the original *l*_*1*_-penalty is replaced, respectively, by the AL penalty (Zou, 2006) and the SCAD penalty (Fan and Li, 2001). Finally, Zhao et al. (2012) introduced the estimating method, high-dimensional undirected graph estimation (Huge) and a companion R package huge, which integrates many functions for estimating graphical models such as semiparametric transformation, graph estimation, and model selection.

To find the optimal regularization parameter (which controls the sparsity of the graph) and hence the optimal sparse undirected graph, several selection criteria have been proposed. For example, Akaike (1974) proposed the well-known Akaike information criterion (AIC) to select the potential optimal model out of a collection of candidate models. Schwarz (1978) proposed the Bayesian information criterion (BIC), which also selects the potential optimal model out of a collection of candidate models but penalizes more for extra parameters in the model than AIC. It has been shown that the BIC consistently selects the optimal model. Fan et al. (2009) introduced a *K*-fold cross-validation score to conduct selection of the optimal graphical model. Moreover, the Huge method is accompanied by three selection criteria: the rotation information criterion (ric) (Lysen, 2009), the extended BIC (ebic) (Foygel and Drton, 2010), and the stability approach for regularization selection (stars) (Liu et al. 2010), all of which are provided by the function huge.select() of the package huge (Zhao et al., 2012).

While some of these methods and selection criteria for estimating sparse graphical models have been introduced to the neuroimaging community (Chouinard et al., 2017; Cribben and Fiecas, 2016; Cribben et al., 2012; Cummine et al., 2016; Grosenick et al., 2013; Pircalabelu et al., 2015; Smith et al., 2011; Westbury et al., 2016), there has never been a thorough validation and comparison study on their practical use in general, nor in particular for estimating FC networks. Specifically, it is not known which combination of estimating method and selection criterion has optimal performance under different experimental settings.

In this article, we have two main objectives. First, we compare several sparse graph estimation procedures and several selection criteria mentioned above under various simulation settings, such as different dimensions or sample sizes, different types of data, and different sparsity levels of the true model structures to find the optimal estimating methods and selection criteria combinations. We discuss in detail the superiority and deficiency of each combination. Our evaluation is aimed at the performance of each combination, which is an estimation method with a selection criterion. We compare their ability to correctly detect the existing network connection, their ability to produce sparse estimates, and their robustness against the violation of some assumptions for the estimation method or the selection criterion. We find that some estimation methods and selection criteria are not effective and always provide undesirable results, but some others can provide satisfactory results, even when some of the assumptions of the method are not met. Our focus is on FC networks and we consider sparse estimation methods because a sparse network structure supports the idea of economic brain organization (Bullmore and Sporns, 2009). Second, we discuss the impact of autocorrelation/whitening on the estimation of FC networks. To this end, we compare the performance of the sparse graph estimation procedures in combination with the selection criteria to independent multivariate normal (MVN) data and to MVN data with an auto-correlation structure. This comparison allows us to make conclusions about the effect autocorrelation in our data has on the estimated FC networks. After our simulation study, we also apply some of the combinations, which are more likely to provide superior estimates to some real fMRI data, to see how they perform in the real world. Our results question the validity of a number of fMRI studies where inferior graphical model techniques have been used to estimate brain networks.

Although the main focus of this work is on estimating methods that estimate static FC where the time series data from each brain region is stationary, the methods can be easily incorporated into an algorithm for estimating dynamic FC via a sliding window or for estimating FC change points in a similar vein to Cribben et al. (2013, 2012).

The rest of this work is organized as follows. The Materials and Methods and Selection Criteria sections explain the theoretical background and features of the estimating methods and selection criteria. The Simulations section is devoted to the simulations, the fMRI data section describes the resting-state fMRI data, and the results from these are discussed in the Results section. In the Discussion section, we discuss the strengths and weaknesses of each combination of estimating methods and selection criteria and some of the parameter choices in the methods. Finally, the Conclusion section provides a set of conclusions based on the results.

## Materials and Methods

### Notation

In this section, we introduce the required notation. Consider a *p* dimensional data set (e.g., time series from several brain regions), **X** = (*X*_1_, *X*_2_, …, *X*_*p*_) ^[*T*]^, with mean vector ***μ*** and a covariance matrix *Σ*_*p* × *p*_ = (*σ*_*ij*_)_1≤*i, j*≤*p*_. Let

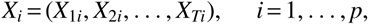

where *T* is the sample size. The (*i, j*) th entry *σ*_*ij*_ of a covariance matrix *Σ*_*p* ×*p*_ is the covariance between *X*_*i*_ and *X*_*j*_, where

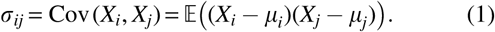

The (*i, j*) th entry of the standardized covariance matrix is the correlation coefficient between *X*_*i*_ and *X*_*j*_. The sample co-variance matrix *S*_*p* × *p*_ = (*s*_*ij*_)_1≤*i, j*≤*p*_ is an empirical statistic calculated from a sample data set whose (*i, j*) th entry *s*_*ij*_ is the sample covariance between the set of observations of *X*_*i*_ and *X*_*j*_ and is estimated using

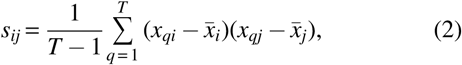

where *T* is the sample size of *X*_*i*_ and *X*_*j*_. The precision matrix *Ω* = (*ω*_*ij*_)_1≤*i, j*≤*p*_ is the inverse of the covariance matrix *Σ* and the (*i, j*) th entry of the standardized precision matrix is the partial correlation between *X*_*i*_ and *X*_*j*_. The estimate of the precision matrix is denoted 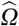.

The precision matrix can also be represented by an undirected graph. Within this framework, graphical models display the dependency structure of a *p* dimensional data set **X** using a graph G. Graphs are mathematical structures that can be used to model pairwise relationships between variables. Let *G* : = (*V, E*) denote a *p*-node undirected simple graph, where *V* : = {1, …, *p*}and *E* : ={(*i, j*), 1 *i* ≤ *j*≤*p*} are the collections of vertices and edges, respectively. The vertices (or nodes) represent a collection of random variables and the edges represent the dependence among these random variables. From a practical point of view, the edge connectivity in a graphical model is the quantity of interest and importance, since it offers an intuitive and effective way of reflecting the underlying network and interplay of the node variables. In this article, we focus exclusively on undirected graphs that do not infer the directionality of dependence or FC between the brain regions. In this study, if the (*i, j*) th component of the precision matrix *Ω* = *Σ*^-1^ is zero, then variables *X*_*i*_ and *X*_*j*_ are said to be conditionally independent, given the other variables, and no edge is included in the graph between the variables. In other words, each entry of the standardization of a precision matrix is a partial correlation coefficient of the corresponding random variables, quantifying their dependence with the influence from all other variables removed. It has been shown that the precision matrix or partial correlations obtain high sensitivity to network connection detection on good-quality fMRI data (Smith et al., 2011). Thus, the central theme of this work is the estimation of a sparse (standardized) precision matrix or a sparse undirected graphical model where some of the elements of the precision matrix (or edges in the undirected graphs) are set exactly to zero. A sparse graphical model is usually preferred in practice for its simplicity and ease of interpretation.

Finally, we do not distinguish between the terms “network” and “graph” in this article and we will use them interchangeably throughout.

### Estimating methods

In this work, our objective is to find the best method for precision matrix estimation in the context of FC network estimation. We consider eight estimating methods: sample CM **C**_*S*_, sample PCM **P**_*S*_ (these two are simply used as reference methods), graphical lasso (glasso), DP glasso, BG, graphical lasso with AL penalties, graphical lasso with SCAD penalties, and Huge.

The glasso, DP glasso, BG, AL, and SCAD methods estimate sparse precision matrices and are based on penalizing the log-likelihood of an MVN distribution. Specifically, they add various weighted *l*_*1*_-penalties to the log-likelihood formula. The estimate of the precision matrix, *Ω*, is the solution to the following formula:

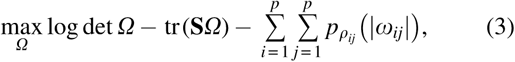

where *tr* denotes the trace of a matrix, which is the sum of all elements on the main diagonal, *Ω* is any positive definite matrix, S is the sample covariance matrix, det *Ω* is the determinant of the matrix *Ω, ω* _*ij*_ the elements of the matrix *Ω*, and 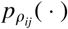 is the penalty function with *ρ*_*ij*_ being the corresponding regularization parameter that controls the sparsity level. We now introduce the estimating methods.

### Sample CM

The (*i, j*) th element, *r*_*ij*_, of the sample CM **C**_*S*_ = (*r*_*ij*_)_*i, j*≤*p*_ is the sample correlation between the *i*th and the *j*th brain regions *X*_*i*_ and *X*_*j*_. It measures the direction and strength of the linear relationship and is estimated using

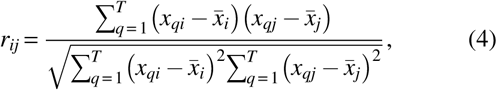

where *T* is the sample size of the time series from brain regions *X*_*i*_ and *X*_*j*_.

### Sample PCM

The (*i, j*) th element, *γ*_*ij*_, of the sample PCM **P**_*S*_ is the sample partial correlation between brain regions, *X*_*i*_ and *X*_*j*_. It measures the relationship between the two brain regions while controlling for the effect from other brain regions, hence providing us with a conditional dependence measure between these two brain regions. The partial correlation is an important measure of dependence when other brain regions are very likely to have effects on *X*_*i*_ and *X*_*j*_. In addition, *γ*_*ij*_ can be estimated from the corresponding elements in the precision matrix *Ω* (Pourahmadi, 2011) using

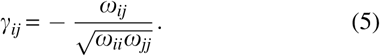

### Graphical lasso

The lasso 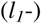 penalty proposed by Tibshirani (1996) has been widely used to estimate sparse undirected graphs. The penalized log-likelihood (2) with the lasso 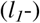 penalty on *Ω* is given by

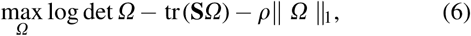

where ‖ *Ω* ‖_1_ denotes the lasso 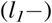 penalty on *Ω* and is the sum of the absolute values of the elements of *Ω*. The 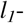 penalty induces sparsity and regularization on the elements of the estimated precision matrix. The tuning parameter *ρ* controls the sparsity of the precision matrix with large values giving rise to a very sparse precision matrix and small values giving rise to a very “full” precision matrix or graph.

Friedman et al. (2007b) developed an efficient algorithm, the blockwise coordinate descent approach (Banerjee et al., 2008), for solving (5). The approach is simple, yet extremely fast, and is named the graphical lasso (glasso). Some elements of 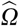 can be shrunk exactly to zero by the glasso algorithm due to the *l*_*1*_-penalty on *Ω*: to maximize (5), *ρ*‖ *Ω* ‖_1_ should be small to make the sum of all absolute values of *Ω* small for a fixed *ρ*, hence some entries of *Ω* are shrunk to zero. The glasso algorithm proceeds by estimating a single row (and column) of *Ω* in each iteration by solving a modified lasso regression.

The R package glasso (Friedman et al., 2007b) can be downloaded to run the above glasso algorithm. The sample covariance matrix, **S**, and the regularization parameter, *ρ*, are two of its inputs. We can also specify the maximum number of iterations of the outer loop (default 10,000), the type of start (starting values for *Σ* and *Ω*, with the default being the cold start **S** + *ρ***I**; another option, the warm start, provides a customized starting value), and a threshold for convergence (default 1*e* - 4), in the package. The glasso package can return 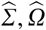 [the maximized value from Eq. (6)], the number of iterations of the outer loop used by the algorithm, and many other outputs.

We denote the estimated precision matrix and covariance matrix from glasso as 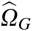 and 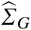, respectively.

### Bootstrap glasso

The BG (Cribben et al., 2013) is a hybridized algorithm of glasso and the bootstrap (resampling scheme), which is inspired by a technique called SS (Meinshausen and Bühlmann, 2010). SS combines subsampling with existing high-dimensional structural selection schemes. Similarly, BG is not a new variable selection technique, but simply aims to enhance existing methods, such as variable selection methods, graphical modeling methods, or cluster analysis methods. In particular, BG does not choose one best model along the whole regularization path, instead, BG bootstraps the original data set a number of times, then keeps structures or variables whose occurrences reach a certain threshold level. More specifically, in graphical model estimation, edges with high selection probabilities remain in the estimate when their selection probabilities are greater than a predefined cut-off 0 < *π*_*thr*_ < 1, otherwise they will be removed. The BG algorithm executes the following four steps for each regularization parameter *q*_*i*_, where *i* may be from 1 to 100:

*Step 1.* Apply glasso to the original data set **X** and obtain the glasso estimate 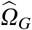.

*Step 2.* Resample (with replacement) **X** *H* times without changing the sample size, obtaining *H* resampled data sets, say 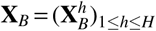.

*Step 3.* Apply glasso to each resampled data set 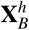 and obtain new glasso estimates, 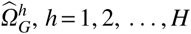.

*Step 4.* For a predefined threshold, 0 < *π*_*thr*_ < 1 (usually set to a value between 0:75 and 0:9), the BG estimate, 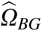, is obtained by setting

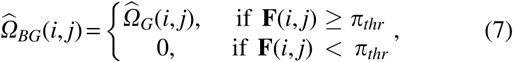

where **F** is the frequency matrix of a nonzero estimate for each element of the matrix *Ω* after resampling *H* times. For example, if **F**(*i, j*) = 0:9, then 90*%* of the glasso estimations on the resampled data are nonzero for *Ω* (*i, j*). Intuitively, 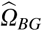 is at least as sparse as 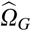, since 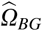 only retains (removes) those nonzero elements of 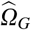, which are estimated as nonzeros (zeros) with high frequency.

### Glasso with AL and SCAD penalties

Two major challenges in estimating sparse precision matrices are (1) constraining *Ω* to be positive definite when optimizing the penalized log-likelihood and (2) minimizing the bias arising from the penalties. For example, it has been shown that glasso, which uses the lasso penalty on *Ω*, is biased (Fan and Li, 2001). To remedy this, the nonconcave SCAD penalty and the AL penalty were proposed by Fan and Li (2001) and Zou (2006), respectively. The AL penalty and the SCAD penalty have the following three desirable properties of an estimator:

1. sparse estimates
2. consistent model selections
3. unbiased estimates for large coefficients

### Adaptive lasso

The AL assigns various weights to each element of *Ω*, where the weights depend on the magnitude of the elements of a consistent estimate of 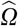. In this study, larger elements of 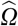 are given smaller weights. Hence, the AL is a properly weighted version of glasso. The penalized log-likelihood with an AL penalty is given by

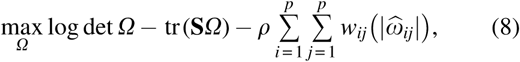

where *w*_*ij*_ is the adaptive weight function (penalty function) and 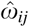 is an estimate for *Ω*(*i, j*). Fan et al. (2009) defined the adaptive weights to be

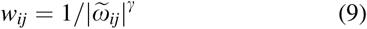

for tuning parameter *γ* > 0, where 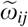 is the (*i, j*) th entry for any consistent estimate 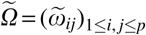. AL has good asymptotic properties, including the property that as the sample size becomes large, the estimate 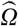 has the same sparsity pattern as *Ω* (Fan and Li, 2001).

The optimal estimate of *Ω* by AL, along a given regularization path, is denoted by 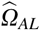.

### Smoothly clipped absolute deviation

The log-likelihood (2) with the SCAD penalty is given by

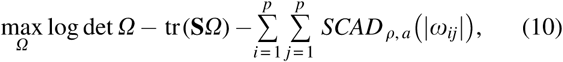

where we use *ρ*_*ij*_ = *ρ* for convenience. Mathematically, the SCAD penalty is symmetric and a quadratic spline on [0, ∞), whose first-order derivative is

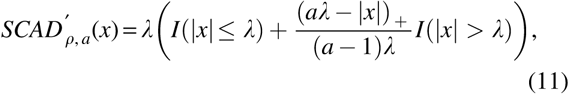

for *x* ≥ 0, where *I* is an indicator function, with *λ* > 0 and *a* > 2 being two tuning parameters. If *a* = ∞, (10) becomes the lasso penalty.

By applying the local linear approximation approach (Zou and Li, 2008) to the SCAD penalty, the original non-concave penalized log-likelihood (2) is transformed into a series of weighted lasso penalized log-likelihood problems, where the weights are controlled by the derivative of the SCAD penalty function. Thus, optimizing the penalized log-likelihood subject to a positive definite *Ω* can be solved iteratively by the efficient glasso algorithm. Consequently, the bias of the penalty is well controlled without losing computational efficiency. Note that in the iterative procedure for SCAD, an estimated zero in 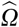 in one step does not necessarily mean it is zero in the next iteration step, whereas for AL, zero estimates will remain zero in each iteration step, and hence, the initial value will always provide denser estimates for AL. The optimal estimate of *Ω* by SCAD, along a given regularization path, is denoted by 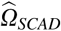.

The AL and SCAD penalties are considered improvements over the glasso as both AL and SCAD can obtain sparse estimates, consistent model selection, and unbiased estimates simultaneously, all of which are not achieved by glasso. However, the simple yet fast glasso algorithm can still be applied to AL and SCAD penalties to estimate sparse *Ω*, as long as the SCAD penalty is locally linearly approximated.

### DP-glasso

As we have already noted, glasso is a popular and efficient algorithm for estimating precision matrices for brain networks (Cribben et al., 2012). However, glasso operates on the dual problem of (5) with the target estimation matrix being the covariance matrix *Σ*, rather than the primal problem itself (the estimation of *Ω*), which results in many undesirable outcomes. Consequently, Mazumder and Hastie (2012) proposed a new method called DP-glasso, which directly solves the primal problem by block coordinate descent, whose optimized matrix is the precision matrix *Ω* and not the covariance matrix. Several advantages arise from this new algorithm, including computational speed. In addition, an R package called dpglasso allows us to implement this algorithm and compare it with the other existing methods.

In the glasso package, the input regularization parameter *ρ* can either be a scalar or a matrix, and thus, the AL and SCAD methods can be implemented easily in it. However, unlike glasso, the regularization parameter required by dpglasso package has to be a scalar, which means the AL and SCAD cannot take advantage of the DP-glasso algorithm directly. Therefore, to take advantage of the DP-glasso algorithm, we use the DP-glasso estimated precision matrices as initial values for AL and SCAD, while the glasso() function is used in the inner steps of AL and SCAD. This leads to another two algorithms denoted as DP-AL and DP-SCAD. For BG, dpglasso() is applied in every step in the procedure, resulting in another method named DP-BG.

We denote the precision matrix estimated by DP-glasso as 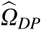. Unlike glasso, DP-glasso returns a sparse and positive definite estimated precision matrix. Similar computational time is consumed by the glasso and DP-glasso algorithms for large values of the regularization parameter. For smaller values of the regularization parameters, DP-glasso is more efficient.

### High-dimensional undirected graph estimation

Huge’s main objective is to estimate high-dimensional undirected graphs while incorporating the many suggestions from Friedman et al. (2007a, b, 2010). Huge integrates many functions, such as data generating, graph estimation, model selection, estimation visualization, and more. Specifically, this merges many up-to-date proposals and results, such as nonparanormal transformation (for non-normal data) and correlation screening approaches for estimating graphs (Fan and Lv, 2008; Liu et al., 2009), as well as the stars approach for stability-based graphical model selection (Liu et al., 2010). In addition, two screening rules are available, lossless screening (Witten et al., 2011) and lossy screening. Huge is available to download as an R package, called “huge.”

Three graph estimation methods are available in huge(): the Meinshausen–Bühlmann (mb) approximation (Meinshausen and Bühlmann, 2010), the graphical lasso (glasso) algorithm (Banerjee et al., 2008; Friedman et al., 2007b), and the thresholded correlation graph estimation method (ct). The speed of the first two methods can be increased further by using the lossy screening rule, which preselects nearby brain regions to the regions of interest (ROIs) using the thresholded correlation before graph estimation [via the scr argument in huge()]. The third method is a variation that is computational efficient and has been widely used in biomedical research (Langfelder and Horvath, 2008).

Finally, the function huge.npn() applies the nonparanormal method in Liu et al. (2009) for estimating a semiparametric Gaussian copula model by truncated normal or normal score. It transforms **X** to a Gaussian distribution to help relax the normality assumption. The optimal precision matrix 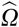 estimated by huge, along the given regularization parameter path, is denoted by 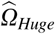.

## Selection Criteria

The penalized log-likelihood methods discussed above contain a regularization parameter that controls the sparsity of the precision matrix. Typically, we estimate the precision matrix along a path of regularization parameters; however, the optimal value of the regularization parameter is unknown. Hence, we consider several selection criteria to choose the optimal regularization parameter among a set of possible values. The selection criteria AIC, BIC, and fivefold Cross-Validation are applied in conjunction with the estimating methods glasso, BG, AL, SCAD, DP-glasso, DP-BG, DP-AL, and DP-SCAD. For the Huge method, we apply the following criteria: ric (Lysen, 2009), ebic (Foygel and Drton, 2010), and stars (Liu et al., 2010), which are embedded in the huge package.

### Akaike information criterion

AIC is a model selection criterion based on combining the likelihood function with a penalty term that guards against overfitting. Hence, it balances the dual needs of adequate model fit and model parsimony. The formula for AIC is

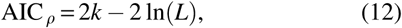

where *k* is the number of nonzero elements in 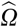 and *L* is the likelihood function, namely (5) without the penalized term 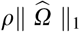. The regularization parameter corresponding to the minimum AIC value gives rise to the best estimation selected by AIC. AIC does not consistently select regression models, that is, if the true model is among the estimating regression models, the probability of selecting the true model by AIC does not approach 1 as *n* → ∞. Hereafter, let *ρ*_*a*_ denote the optimal regularization parameter selected by AIC.

### Bayesian information criterion

BIC is also a model selection criterion based on combining the likelihood function but penalizes more for extra parameters in the model than AIC. The formula for BIC is

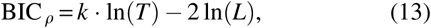

where *T* is the sample size, *k* is the number of nonzero elements in 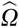, and *L* is the likelihood function (Schwarz, 1978). The *ρ* value that gives rise to the minimum BIC value is optimal. An underlying assumption of BIC is that the observations are independent and identically distributed (Schwarz, 1978). If the observations are not i.i.d, then the effective sample size is not *T* and the formula must be adjusted. BIC consistently selects regression models unlike AIC (Nishii, 1984). For linear regression models, the model chosen by BIC is either the same or a simpler version than that chosen by AIC, due to the heavier penalty (Shao, 1993). Hereafter, let *ρ*_*b*_ denote the optimal regularization parameter selected by BIC.

### *K*-fold cross-validation

The *K*-fold cross-validation score (Fan et al. 2009) is given by

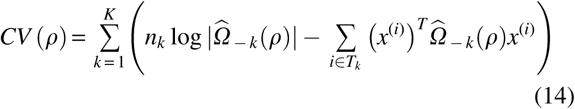

where *n*_*k*_ is the size of *k*th fold *T*_*k*_ and 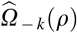 is the estimate of the precision matrix based on the sample 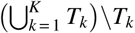 (the training data). The *ρ* that provides the minimum cross-validation (CV) value is the best regularization parameter. In our work, we use fivefold CV to choose the optimal regularization parameter. As the sample size grows larger, minimizing the AIC is equivalent to minimizing the CV for any model, not just linear models (Stone, 1974). Generally, CV does not consistently select models (Shao, 1993). CV performs poorly with high-dimensional data, sometimes dramatically (Meinshausen and Bühlmann, 2010). Hereafter, let *ρ*_*c*_ denote the optimal regularization parameter selected by CV.

### Selection criteria for Huge

Huge provides three selection criteria for choosing the best estimate of the precision matrix: ric, ebic, and stars. ric is a newly developed and very efficient selection criterion. It directly chooses the best regularization parameter *ρ* based on random rotations rather than finding the best *ρ* over the whole regularization path using time-consuming techniques such as cross-validation or subsampling. More specifically, the brain regions are randomly rotated several times so that the minimum *ρ*, which generates all zeros estimated by using the rotated data, will be selected. Thus far, there has been no theoretical proof for consistent selection by ric. In addition, ric suffers from overselection and frequently from underselection. Hence, Zhao et al. (2012) stated that if false-negative levels (few missing selections) are expected, then the number of rotations for ric should be increased, or the selection criterion stars should be applied. ric is available for all three estimation methods provided by the R package, huge.

We denote ebic as BIC _*γ*_, where 0 ≤ *γ* ≤ 1 is called the ebic parameter. The original BIC is equivalent to BIC _0_ (i.e., *γ* = 0). *γ* = 0:5 is the default setting in huge.select(). It has been shown in Chen and Chen (2008) that BIC _1_ is consistent as long as the dimension *p* (or the number of brain regions) does not grow exponentially with the sample size. In huge, we can only use ebic selection criterion for the glasso method.

stars selects the optimal precision matrix in a similar manner to the subsampling procedure discussed above. Hence, it is not computationally efficient. Under certain conditions, stars is shown to be partially consistent but suffers from the problem of overselection in estimating Gaussian graphical models while its performance also depends on the regularization parameters used. Moreover, stars can be used for all three estimation methods in huge, which are the Meinshausen– Bühlmann approximation (mb), glasso (glasso), and thresholded correlation estimation (ct).

## Simulations

The estimating methods and selection criteria described above have many theoretical results. However, there has not been an extensive simulation study to compare the estimating methods in combination with the selection criteria for exploring brain networks. In this work, we compare their performance using data generated with different dimensions, sample sizes, and underlying distributions. As neuroimaging data (fMRI, EEG, MEG, ECoG) are inherently autocorrelated, we also consider this data structure. In what follows, we describe the setup of our data and introduce evaluation criteria for comparing the combination of estimating methods and selection criteria.

### Simulation setup

Let *M*_*q*_ denote the *q*th estimating method, *q* = 1, …, 11, and let *C*_AIC_, *C*_BIC_, *C*_CV_, *C*_ric_, *C*_ebic_, and *C*_stars_ denote the selection criteria AIC, BIC, CV, ric, ebic, and stars, respectively. Also, let 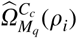 denote the precision matrix estimated by method *M*_*q*_ using selection criteria *C*_*c*_ when the regularization parameter is *ρ*_*i*_. In addition, let *ρ* (*M*_*q*_ * *C*_*c*_) denote the best regularization parameter along the path *ρ*_1_, …, *ρ*_100_ for estimation method *M*_*q*_ and selection criteria *C*_*c*_. The resulting estimated precision matrix 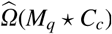 is the best estimate using estimating method *M*_*q*_ and selection criterion *C*_*c*_. The setup of our simulation study is as follows:

*Step 1.* Simulate a data set **X** (which is of dimension *T* × *p*, where *T* and *p* represent the number of time points and brain regions/voxels, respectively).

*Step 2.* Fix 100 equally spaced regularization parameters *ρ* ∈ [0:01, 1] with *ρ*_*i*_ = *i* × 0:01, *i* = 1, …, 100.

*Step 3.* Apply each estimating method *M*_*q*_ to **X** and obtain 100 estimated precision matrices

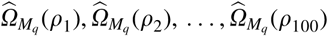

corresponding to the 100 regularization parameters *ρ*_1_, …, *ρ*_100_.

*Step 4.* Choose the estimate from 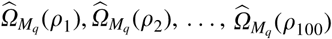 that minimizes the selection criteria formula.

*Step 5.* For each method *M*_*q*_ and selection criterion *C*_*c*_, repeat the above procedure *L* times. This provides *L* best estimated precision matrices for each combination of estimating method and selection criterion.

*Step 6.* Use the *L* estimated precision matrices to evaluate the performance of the estimating methods in combination with the selection criteria.

### Evaluation standards

The estimated precision matrices are compared to the true precision matrices using three evaluation standards. The first two standards, called *True Positive* (TP) and *True Negative* (TN), are numeric and measure the estimation accuracy. The third standard, named the *Average Sparsity Pattern* (ASP) plot, is a plot that provides a visual depiction of the sparsity levels of the estimated matrices.

Our definition of TP is

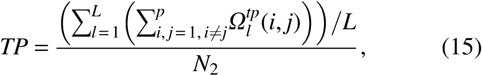

where *L* is the number of simulations and *N*_*2*_ is the number of off-diagonal nonzero entries in the true precision matrix. The TP matrix 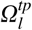 for the *l*th simulation is defined by

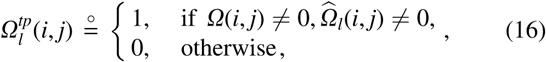

where 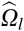 is the estimated precision matrix in the *l*th repetition. If both the true and estimated entry in the (*i, j*) th position in the precision matrix are nonzeros, the (*i, j*) th entry of the TP matrix, 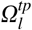, is equal to 1, otherwise it is 0. Thus 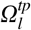 records whether each nonzero true entry is successfully detected in each simulation. TP is a number between 0 and 1 and reflects the average proportion each combination estimates nonzero entries correctly. The larger the TP, the superior the combination method. TP = 1 indicates that all the nonzero entries in O are estimated as nonzeros across all simulations. TP = 0 indicates that none of the true nonzero entries in O was estimated correctly, that is, all of the nonzero entries were estimated as zeros across all simulations.

The second standard TN is defined similarly by

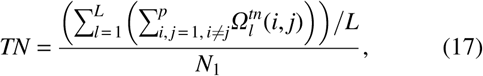

where *N*_*1*_ is the number of off-diagonal zero entries in the true precision matrix, *Ω*. The TN matrix 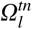 for the *l*th simulation is defined by

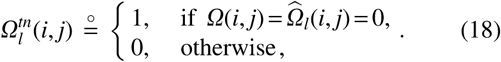

Similar to the TP matrix, the TN matrix marks down whether each zero entry in *Ω* is successfully estimated to be zero. TN tells us how often zero entries are estimated as zeros. TN is also a numeric value between 0 and 1, with higher values indicating a superior combination method. Typically, larger TN means sparser estimated graphs. We did not consider the F-1 score as the TP and TN matrices provide more detail on the estimation of nonzeros and zeros.

The ASP plot is obtained by plotting the ASP matrix, ASP _*p* × *p*_. For each *i, j* = 1, …, *p*, ASP(*i, j*) is defined by

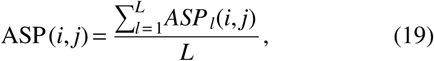

with the matrix ASP _*l*_(*i, j*) of the *l*th simulation defined by

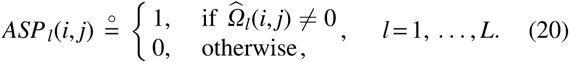

Hence, the matrix ASP _*p* × *p*_ shows the percentage of times each element of the precision matrix is estimated as nonzero across all simulations. The larger the ASP value, the darker the corresponding rectangular area in the plot. Hence, the more dense the estimated precision matrices, the darker the ASP plot. This is an intuitive and clear way to show the overall sparsity levels of the precision estimates.

### Simulation settings

#### BG parameters

For BG, we fix the number of resamples to 50 and the threshold value to *π*_*thr*_ = 0:9.

#### AL parameters

For AL, we use *γ* = 0:5 in the penalty (7) as there are no obvious differences among estimates using different *γ* values (Fan et al., 2009). As the AL requires consistent estimates, we can use the precision matrix 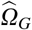 estimated by the glasso for the initial value of its penalty. That is, we set 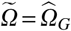 and 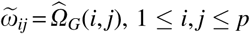 in the AL penalty (7). Fan et al. (2009) noted that we can use the inverse sample covariance matrix **S**^- 1^ for 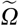 in low-dimensional cases (*p* < *T*) and 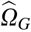 in the moderate-dimensional cases (*p* ≥ *T*). However, **S**^- 1^ might be inconsistent if *p* increases at the same rate as *T*. The requirement for a consistent initial value is one of the drawbacks of AL. The R package glasso is convenient for implementing the AL algorithm. We first calculate the AL penalty matrix for each *ρ* and then apply this penalty matrix using the rho argument in the package glasso to conduct the estimations by AL.

#### SCAD parameters

To minimize the Bayes risk, Fan and Li (2001) recommended using *a* = 3:7 in (10), which we used in our simulations. The precision matrix 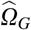 estimated by the glasso is used as the initial value for SCAD. The precision matrix estimation by SCAD can also take the advantage of the package glasso. Similar to AL, we first calculate the SCAD penalty matrix for each *ρ*, then set this as the rho argument in the package glasso. We then choose the best *ρ* that minimizes a selection criterion and use this optimal *ρ* to iteratively obtain a new estimated precision matrix. We stop the iterative procedure when the difference between the sum of the absolute values between the two estimated precision matrices is less than a threshold, which we set to be 1*e* - 04.

#### Huge parameters

In the function huge(), we choose glasso for the method argument and set *scr* = FALSE, so the lossy screening rule is not applied to preselect the neighborhood before graph estimation. After running the function huge(), an object with class S3 is returned and contains values, including icov, a list of *p* × *p* estimated precision matrices corresponding to each regularization parameter, and loglik, a vector with the same length as *ρ*, which contains the log-likelihood values along the regularization path. To implement huge.select(), the S3 class object from huge() is the first required argument. Accordingly, huge.select() picks the best estimate along the whole regularization path. All three Huge criteria (ric, ebic, stars) provided by huge.select() are applied in our simulation study.

#### Data settings

The number of simulations, *L*, is 100. Specifically, we apply each combination of estimating method and selection criterion to the same 100 different data sets for various data types and then observe how the combinations perform on average over the 100 simulations. Hence, the differences between the results only arise from the estimating methods and selection criteria themselves.

#### Regularization path settings

In all our simulations, 100 equally spaced regularization parameters *ρ* are used, namely *ρ* ∈ [0:01, 1] and *ρ*_*i*_ = *i* × 0:01, *i* = 1, …, 100. A possible drawback of setting *ρ* [0:01, 1] is that there may exist some smaller *ρ*s that could provide superior estimates than our best estimate. Another drawback is that our regularization parameters are discrete. Hence, it may be the case that some other *ρ*s, which are not a multiple of 0.01 (e.g., 0.233), could produce better estimates. Nevertheless, we believe our regularization parameter choices and working procedures are convincing and the estimating methods are comparable. All the methods are supplied with the same path of *ρ*s and they work under the same level of parameter precision and carry out the same operations. In addition, our working regularization parameters are relatively dense and have a wide range.

#### Simulations

We use simulations to examine the performance of the penalized log-likelihood approaches to estimate the precision matrix. In each example, we first generate a true precision matrix *Ω*, which will be fixed for the whole example. Next, we generate a data set of *T* independent and identically distributed random vectors distributed as N(0, *Ω*^-1^). We name this the MVN data. As voxels and brain regions from fMRI data are inherently autocorrelated, we also simulate MVN data that follow an AR(1) model: *Y*_*t*_ = *ϕ*_1_*Y*_*t* – 1_+ *ε*_*t*_, where *ε*_*t*_∼N(0, 1). We call the MVN data where every brain region (or column of **X**) has an AR1 autocorrelation structure, MVNAR1. The two data types allow us to look into how the autocorrelation structure affects the network estimation. In particular, they allow us to evaluate the deterioration in performance of the estimating methods if prewhitening of the time series data from each brain region is not carried out. In the simulation study, we consider two extremes, completely independent data and strongly autocorrelated data, and hence, we set the autocorrelation parameter equal to 0.8 (*ϕ*_1_ = 0:8).

#### Low-dimensional simulations

As some brain imaging studies consider only a small number of brain regions in their network analysis, we include simulations that are of small dimension. For these cases, we choose *p* = 5 (five brain regions). The two data types are studied using four different sample sizes (*T* = 100, 200, 500, 1000) to observe how the results vary for different sample sizes. The true precision matrix *Ω* we study is of general form and is given by

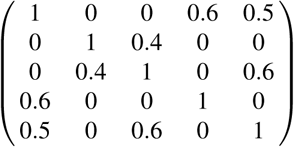

#### Moderate-dimensional simulations

As many neuroimaging studies consider only a moderate number of brain regions in their analysis, our moderate-dimensional case examines *p* = 30 brain regions. In this case, we study three different true precision matrices—the *tridiagonal matrix*, the *exponential decay matrix*, and the *general matrix*—and evaluate how each combination performs. We now detail the schemes for generating these matrices.

For the tridiagonal case, where dependence between the brain regions is high close to the main diagonal and zero everywhere else, the (*i, j*) th element of *Ω* is defined to be

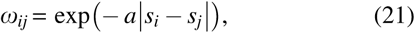

where *a* is a positive constant and *s*_*i*_, *s*_*j*_ are random values such that *s*_1_ < *s*_2_ < … < *s*_*p*_ and

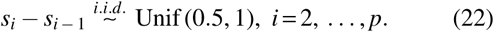

Obviously, a larger *a* value produces smaller off-diagonal elements in *Ω*. In this study, we use the tridiagonal matrix with *a* = 1:7, which results in nonzero entries in the precision matrix close to 0:3.

For the exponential decay case, where dependence between the brain regions decays the further from the main diagonal, no element in the precision matrix *Ω* is exactly zero, but it contains a number of entries close to 0. The (*i, j*) th element of the true exponential decay precision matrix is defined to be

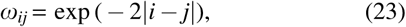

which can be extremely small when |*i j*| is large. Since none of the entries of the true exponential decay matrix *Ω* is exactly zero, we set a threshold of 1*e* − 03 when calculating the TP and TN. Otherwise, the TN will have value NA and the TP will be dramatically small, which will obscure the results. However, the true *Ω* without thresholding is still used to generate the original data set. Also, in the SP plots, a threshold is applied to the true precision matrix, *Ω*.

For the general matrix case, we generate an upper triangular matrix first. Each element in the upper triangular matrix is generated uniformly over [−5, −1] ∪ [1, 5]. Then, the smallest 50% of the entries are set to zero and the remaining nonzeros in the matrix are randomly dispersed. By symmetrizing this upper triangular matrix, we get a matrix with main diagonals equal to 0. We set the (*i, i*) th entry in this symmetric matrix to be a multiple of the sum of the absolute values of the *i*th row elements. In this study, we choose a multiple of 2 to ensure the resulting O is positive definite. Table 1 provides a summary of the entries of the three precision matrices generated as above. Figure 1 shows the sparsity patterns of the three true precision matrices.

**TABLE 1.** SUMMARY OF THE ENTRIES OF THE TRIDIAGONAL MATRIX WITH *a* = 1:7, THE EXPONENTIAL DECAY MATRIX AND THE ABSOLUTE VALUES OF THE GENERAL MATRIX FOR THE MODERATE-DIMENSIONAL MULTIVARIATE NORMAL DATA AND THE MULTIVARIATE NORMAL DATA WITH AN AR1 AUTOCORRELATION STRUCTURE

|  | *Smallest* | *Largest* | *0.1 to 1* | *0.01 to 0.1* | *Less than 0.01* |
| --- | --- | --- | --- | --- | --- |
| Tridiagonal | 0.191 (except 0) | 0.425 | 0.067 | 0 | 0.933 |
| Exponential decay | $6.470 \cdot e - 26$ | 0.59 | 0.067 | 0.064 | 0.869 |
| General | 0.019 | 0.621 | 0.057 | 0.443 | 0.499 |

**FIG. 1.**
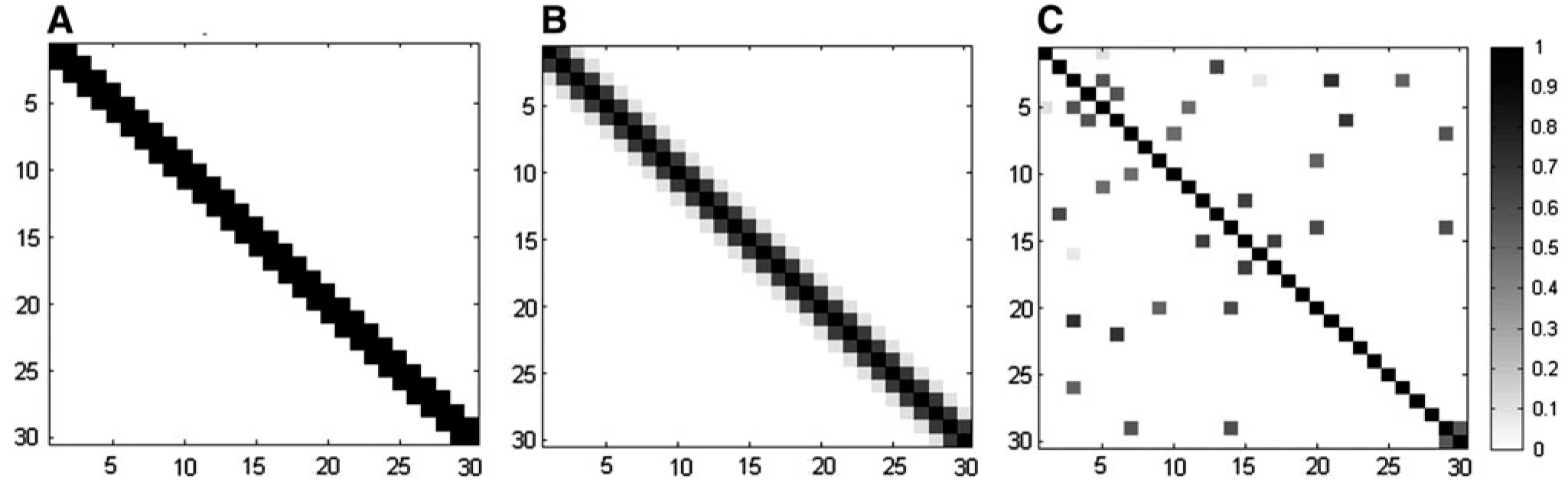
The true sparsity patterns of **(A)** the tridiagonal matrix with *a* = 1:7, **(B)** the exponential decay matrix, and **(C)** the general matrix.

## fMRI Data

We applied the combination of estimating methods and selection criteria to a resting-state fMRI data set, as described in Habeck et al. (2012). Participants (*n* = 45) were instructed to rest in the scanner for 9.5 min, with the instruction to keep their eyes open for the duration of the scan. Functional images were acquired using a 3.0 Tesla magnetic resonance scanner (Philips) using a field echo echo-planar imaging sequence (TE/TR = 20 ms/2000 ms; flip angle = 72; 112 × 112 matrix; in-plane voxel size = 2.0 × 2.0 mm; slice thickness = 3.0 mm [no gap]; 37 transverse slices per volume). In addition, a T1-weighted turbo field echo high-resolution image was also acquired (TE/TR = 2.98 ms/6.57 ms; flip angle = 8; 256 × 256 matrix; in-plane voxel size = 1.0 × 1.0 mm; slice thickness = 1.0 mm [no gap]; 165 slices). The individual time series data were bandpass filtered between 0.009 and 0.08 Hz, motion corrected, and coregistered to the structural data, with a subsequent spatial normalization to the Montreal Neurological Institute (MNI) template. The voxel time courses at white matter and cerebrospinal fluid (CSF) locations were submitted to a principal components analysis and, together with the motion parameters, we used all components with an eigenvalue strictly >1 as independent variables in a subsequent nuisance regression. Each voxel’s time series was residualized with respect to those independent variables, that is, it was regressed against the independent variables, and the model prediction was subtracted from the time series voxel to form a residual time series for each subject at each voxel location. The residual time series images were then smoothed with an isotropic Gaussian kernel (full width at half maximum [FWHM] = 6 mm). We applied the Anatomical Automatic Labeling (Tzourio-Mazoyer et al., 2002) atlas to the adjusted voxelwise time series and produced time series for 31 ROIs for each subject by averaging the voxel time series within the ROIs. The 31 ROIs contained 8 regions from the attentional network (frontal superior medial L, angular L, angular R, temporal middle L, temporal mid R, thalamus L, cerebellum crus1 L, cerebellum crus1 R), 2 regions from the visual network (temporal superior L, temporal superior R), 3 regions from the sensorimotor network (postcentral L, post-central R, supplementary motor area R), 7 regions from the salience network (cingulum anterior L, frontal mid L, frontal middle R, insula L, insula R, supramarginal L, supramarginal R), 9 regions from the default mode network (precentral L, pre-central R, parietal superior L, occipital superior R, parietal inferior L, parietal inferior R, temporal inferior L, temporal inferior R, cingulum posterior L), and 2 regions from the auditory network (calcarine L, calcarine R). We chose these networks because an increasing number of pathologic conditions appear to be reflected in the FC between these particular brain regions and we wanted the number of ROIs in the fMRI data to match the simulation settings. In total, each ROI time series is made up of 285 time points (9.5 min with TR = 2).

## Results

### MVN data

#### Low-dimensional cases

Table 2 displays the TP for the MVN data set with dimension *p* = 5 for four different sample sizes. It is clear that large nonzero entries of the precision matrix are easy to estimate correctly, especially when the number of time points, *T*, is large, with less than 1% missed detections across all the combinations of estimating methods and selection criteria. Only SCAD provides zero estimates for true nonzero entries. All the selection criteria perform similarly in estimating nonzeros. In addition, the newly proposed DP-glasso algorithm does not show obvious improvements over glasso, AL, SCAD, and BG. For low-dimensional data, *T* = 100 is sufficiently large to estimate large-value non-zero entries. Moreover, increasing the number of time points does not appear to notably improve the results.

**TABLE 2.** THE TRUE POSITIVE FOR THE MULTIVARIATE NORMAL (*p* = 5) DATA FOR FOUR DIFFERENT SAMPLE SIZES

| | $T = 100$ | | | $T = 200$ | | | $T = 500$ | | | $T = 500$ | | |
| --- | --- | --- | --- | --- | --- | --- | --- | --- | --- | --- | --- | --- |
|  | TP |  |  | TP |  |  | TP |  |  | TP |  |  |
|  | AIC | BIC | CV | AIC | BIC | CV | AIC | BIC | CV | AIC | BIC | CV |
| CM | 1 |  |  | 1 |  |  | 1 |  |  | 1 |  |  |
| PCM | 1 |  |  | 1 |  |  | 1 |  |  | 1 |  |  |
| Glasso | 1 | 1 | 1 | 1 | 1 | 1 | 1 | 1 | 1 | 1 | 1 | 1 |
| AL | 1 | 1 | 1 | 1 | 1 | 1 | 1 | 1 | 1 | 1 | 1 | 1 |
| SCAD | 0.999 | 1 | 0.993 | 0.995 | 0.996 | 0.999 | 1 | 1 | 1 | 1 | 1 | 1 |
| DP |  |  |  |  |  |  |  |  |  |  |  |  |
| Glasso | 1 | 1 | 1 | 1 | 1 | 1 | 1 | 1 | 1 | 1 | 1 | 1 |
| AL | 1 | 1 | 1 | 1 | 1 | 1 | 1 | 1 | 1 | 1 | 1 | 1 |
| SCAD | 0.999 | 0.998 | 0.998 | 0.996 | 0.99625 | 0.999 | 1 | 1 | 1 | 1 | 1 | 1 |
| BG |  |  |  |  |  |  |  |  |  |  |  |  |
| $H = 50$ | 1 | 1 | 1 | 1 | 1 | 1 | 1 | 1 | 1 | 1 | 1 | 1 |
| DP-BG | 1 | 1 | 1 | 1 | 1 | 1 | 1 | 1 | 1 | 1 | 1 | 1 |
| Huge |  |  |  |  |  |  |  |  |  |  |  |  |
| Ric | 1 |  |  | 1 |  |  | 1 |  |  | 1 |  |  |
| Stars | 1 |  |  | 1 |  |  | 1 |  |  | 1 |  |  |
| Ebic | 1 |  |  | 1 |  |  | 1 |  |  | 1 |  |  |

AIC, Akaike information criterion; AL, adaptive lasso; BG, bootstrap graphical lasso; BIC, Bayesian information criterion; CM, correlation matrix; CV, cross-validation; ebic, extended BIC; huge, high-dimensional undirected graph estimation; PCM, partial CM; ric, rotation information criterion; SCAD, smoothly clipped absolute deviation; stars, stability approach for regularization selection; TP, true positive.

Table 3 presents the TN for the MVN data with dimension *p* = 5 for the four different sample sizes. It is clear that the zero entries of the true precision matrix O are more difficult to estimate than the nonzero entries, even for low-dimensional data with a large number of time points. In general, the results indicate that increasing *T* enhances the correct estimation of zeros. SCAD is the best method for detecting zero entries, especially when combined with the selection criterion, BIC. AL’s and BG’s performance are marginally inferior to SCAD. More specifically, in combination with AIC and BIC, BG estimates more zeros than AL, while AL outperforms BG when combined with CV. Glasso’s performance is always inferior to AL, BG, and SCAD at capturing zeros. Again, as in the TP case, the DP-glasso algorithm appears to have no obvious improvements over glasso, AL, SCAD, and BG. Generally, all estimating methods improve as *T* increases.

**TABLE 3.** THE TRUE NEGATIVE FOR THE MULTIVARIATE NORMAL (*p* = 5) DATA FOR FOUR DIFFERENT SAMPLE SIZES

|  | T = 100 |  |  | T = 200 |  |  | T = 500 |  |  | T = 1000 |  |  |
| --- | --- | --- | --- | --- | --- | --- | --- | --- | --- | --- | --- | --- |
|  | TN |  |  | TN |  |  | TN |  |  | TN |  |  |
|  | AIC | BIC | CV | AIC | BIC | CV | AIC | BIC | CV | AIC | BIC | CV |
| CM | 0 |  |  | 0 |  |  | 0 |  |  | 0 |  |  |
| PCM | 0 |  |  | 0 |  |  | 0 |  |  | 0 |  |  |
| Glasso | 0.213 | 0.333 | 0.335 | 0.143 | 0.295 | 0.241 | 0.193 | 0.353 | 0.249 | 0.24 | 0.353 | 0.25 |
| AL | 0.499 | 0.646 | 0.678 | 0.52 | 0.673 | 0.649 | 0.687 | 0.815 | 0.73 | 0.843 | 0.854 | 0.843 |
| SCAD | 0.708 | 0.9 | 0.855 | 0.75 | 0.91 | 0.868 | 0.937 | 0.982 | 0.943 | 0.955 | 0.976 | 0.963 |
| DP |  |  |  |  |  |  |  |  |  |  |  |  |
| Glasso | 0.213 | 0.33 | 0.335 | 0.143 | 0.297 | 0.245 | 0.193 | 0.352 | 0.248 | 0.242 | 0.352 | 0.252 |
| AL | 0.499 | 0.645 | 0.678 | 0.52 | 0.672 | 0.648 | 0.687 | 0.815 | 0.73 | 0.842 | 0.853 | 0.842 |
| SCAD | 0.708 | 0.902 | 0.857 | 0.755 | 0.912 | 0.868 | 0.937 | 0.982 | 0.943 | 0.955 | 0.977 | 0.963 |
| BG |  |  |  |  |  |  |  |  |  |  |  |  |
| H = 50 | 0.672 | 0.858 | 0.558 | 0.595 | 0.836 | 0.496 | 0.7 | 0.877 | 0.695 | 0.828 | 0.893 | 0.83 |
| DP-BG | 0.673 | 0.858 | 0.56 | 0.593 | 0.837 | 0.5 | 0.702 | 0.878 | 0.697 | 0.702 | 0.878 | 0.697 |
| Huge |  |  |  |  |  |  |  |  |  |  |  |  |
| Ric | 0.252 |  |  | 0.17 |  |  | 0.172 |  |  | 0.175 |  |  |
| Stars | 0.243 |  |  | 0.29 |  |  | 0.353 |  |  | 0.374 |  |  |
| Ebic | 0.4 |  |  | 0.338 |  |  | 0.353 |  |  | 0.374 |  |  |

TN, true negative.

Overall, the BIC selection criterion correctly estimates the largest number of zero entries for the glasso, BG, AL, and SCAD estimating methods (as well as for DP-glasso, DP-BG, DP-AL, and DP-SCAD) across essentially all cases, with the only exception being glasso and AL methods when *T* = 100. In this case, CV correctly selects marginally more zeros than BIC. AIC always selects less zero entries than BIC and CV for estimating methods, glasso, AL, and SCAD. However, BG behaves differently in combination with AIC, it correctly selects more zeros than CV when *T* < 1000, but remains inferior to BIC overall.

In general, the ebic criterion correctly estimates the most zeros among the three selection criteria for Huge. However, the Huge * ebic combination is only marginally superior to the glasso estimates and considerably inferior to the SCAD * BIC combination. There is no noticeable improvement in the TN for Huge * ric as the number of time points increases. Also, the TN decreases when the number of time points increases from 100 to 200 for Huge * ric. stars is the only criterion in Huge that the TN increases with the number of time points.

To conclude, in terms of TN, SCAD is the best estimating method compared with glasso, AL, BG, and Huge. We regard AL as the second best approach if we take into consideration the computational time. BG provides competitive estimates to AL, while glasso and Huge are less effective. For selection criteria, BIC selects the best estimates for glasso, BG, AL, and SCAD in most cases. ebic can be considered the best criterion for Huge. SCAD in combination with any of the AIC, BIC, or CV criterion outperforms all other combinations, except that BG*BIC performs better than SCAD*AIC when *T* ≥ 200, which is an indication that the superiority of a selection criterion can remedy the inefficiency of an estimating method.

#### Moderate-dimensional case

Figure 1 shows the sparsity patterns of the three true precision matrices: the tri-diagonal matrix with the constant *a* = 1:7 in (23), the exponential decay matrix, and the general matrix. Figure 2 displays the ASP plots for the sample CMs and the sample PCMs, which are almost identical across all dimensions, *Ω* types, and number of time points: all the entries of both matrices are always nonzero, which is a strong reason that they are not effective for estimating sparse brain networks. Thus, we do not show the ASP plots for CM and PCM hereafter.

**FIG. 2.**
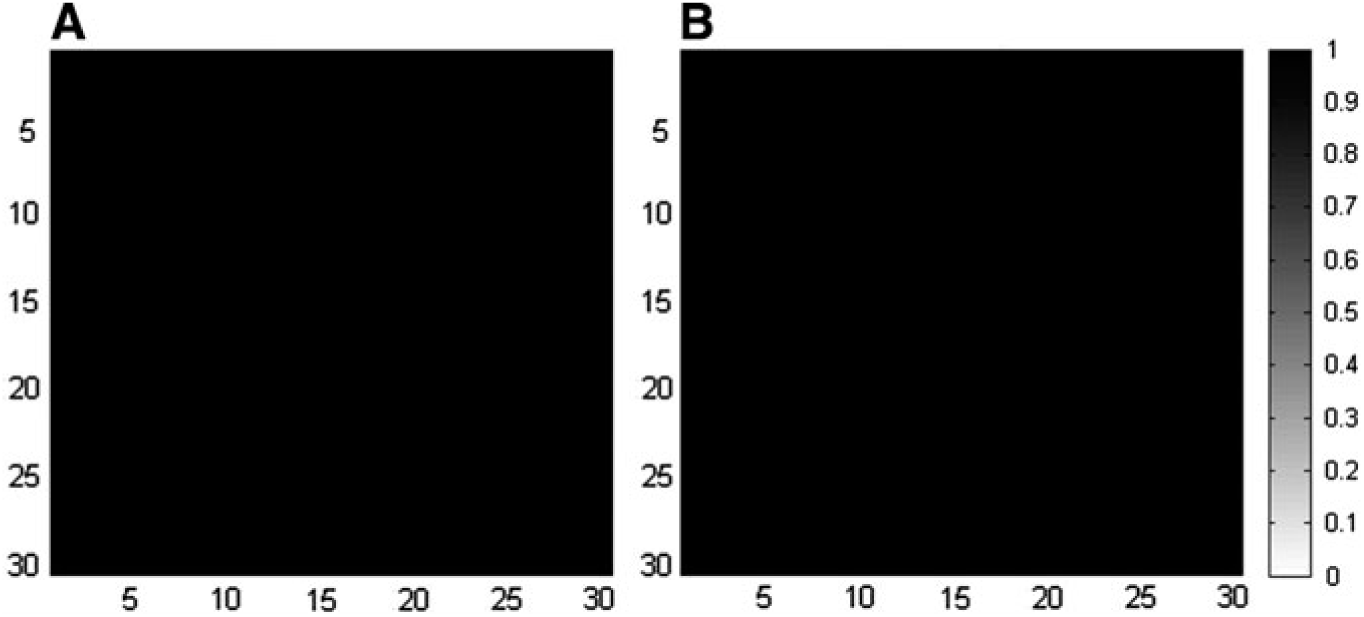
The ASP plots for **(A)** the sample correlation matrix and **(B)** the sample partial correlation matrix. ASP, Average Sparsity Pattern.

Figure 3 contains the ASP plots and Table 4 contains the exact TP and TN for all the combinations of the estimating methods and the selection criteria applied to the moderate-dimensional MVN (*p* = 30) data set generated from the tridiagonal true precision matrix with *a* = 1:7. The table is consistent with the conclusions drawn from the ASP plots. Generally, the larger the TP, the darker the ASP plots and the denser the estimates. Similarly, the larger the TN, the lighter the ASP plots and the sparser the estimates. All the methods perform similarly in terms of TP, but SCAD outperforms the rest in terms of TN. As can be seen from Figure 3, the SCAD*BIC and SCAD*CV combinations produce the sparsest estimates without losing the true graphical structure. For these combinations, almost 90*%* of the zero entries are successfully captured. BG*BIC performs marginally worse than the best SCAD combinations, with a detection rate of the zeros at 85*%*. AL*BIC, glasso*BIC, glasso*CV, BG*CV, and AL*CV result in denser estimates with *80*%* correctly estimated zeros. SCAD is the only estimating method in combination with AIC that does not lead to overly dense estimates. Furthermore, BG*AIC and glasso*AIC estimates are so dense; it becomes more difficult to differentiate the true graphical structure from the noise. For nonzero elements in the precision matrix, all combinations work well in that they seldom estimate the nonzero entries to be zero. This indicates that nonzero entries of a precision matrix that are greater than 0:19 are large enough to be sensitively detected by glasso, BG, AL, and SCAD, where 0:19 is the smallest entry value of the tridiagonal matrix with *a* = 1:7.

**TABLE 4.** THE TRUE POSITIVE AND TRUE NEGATIVE FOR THE TRIDIAGONAL MATRIX WITH *a* = 1:7 FOR THE MULTIVARIATE NORMAL (*p* = 30) DATA

| $\Omega$ | $Tri\ a = 1.7$ | | | | | |
| --- | --- | --- | --- | --- | --- | --- |
| | $TP$ | | | $TN$ | | |
| | $AIC$ | $BIC$ | $CV$ | $AIC$ | $BIC$ | $CV$ |
| CM | 1 |  |  | 0 |  |  |
| PCM | 1 |  |  | 0 |  |  |
| Glasso | 0.995 | 0.958 | 0.967 | 0.172 | 0.783 | 0.741 |
| AL | 0.983 | 0.950 | 0.958 | 0.452 | 0.819 | 0.798 |
| SCAD | 0.961 | 0.915 | 0.925 | 0.661 | 0.908 | 0.897 |
| DP |  |  |  |  |  |  |
| Glasso | 0.995 | 0.958 | 0.967 | 0.172 | 0.783 | 0.741 |
| AL | 0.983 | 0.950 | 0.958 | 0.451 | 0.819 | 0.798 |
| SCAD | 0.958 | 0.915 | 0.925 | 0.684 | 0.908 | 0.897 |
| BG |  |  |  |  |  |  |
| $H = 50$ | 0.989 | 0.932 | 0.962 | 0.267 | 0.855 | 0.756 |
| DP-BG | 0.965 | 0.9 | 0.948 | 0.497 | 0.899 | 0.799 |
| Huge |  |  |  |  |  |  |
| Ric | 0.582 |  |  | 0.998 |  |  |
| Stars | 0.997 |  |  | 0.116 |  |  |
| Ebic | 0 |  |  | 1 |  |  |

**FIG. 3.**
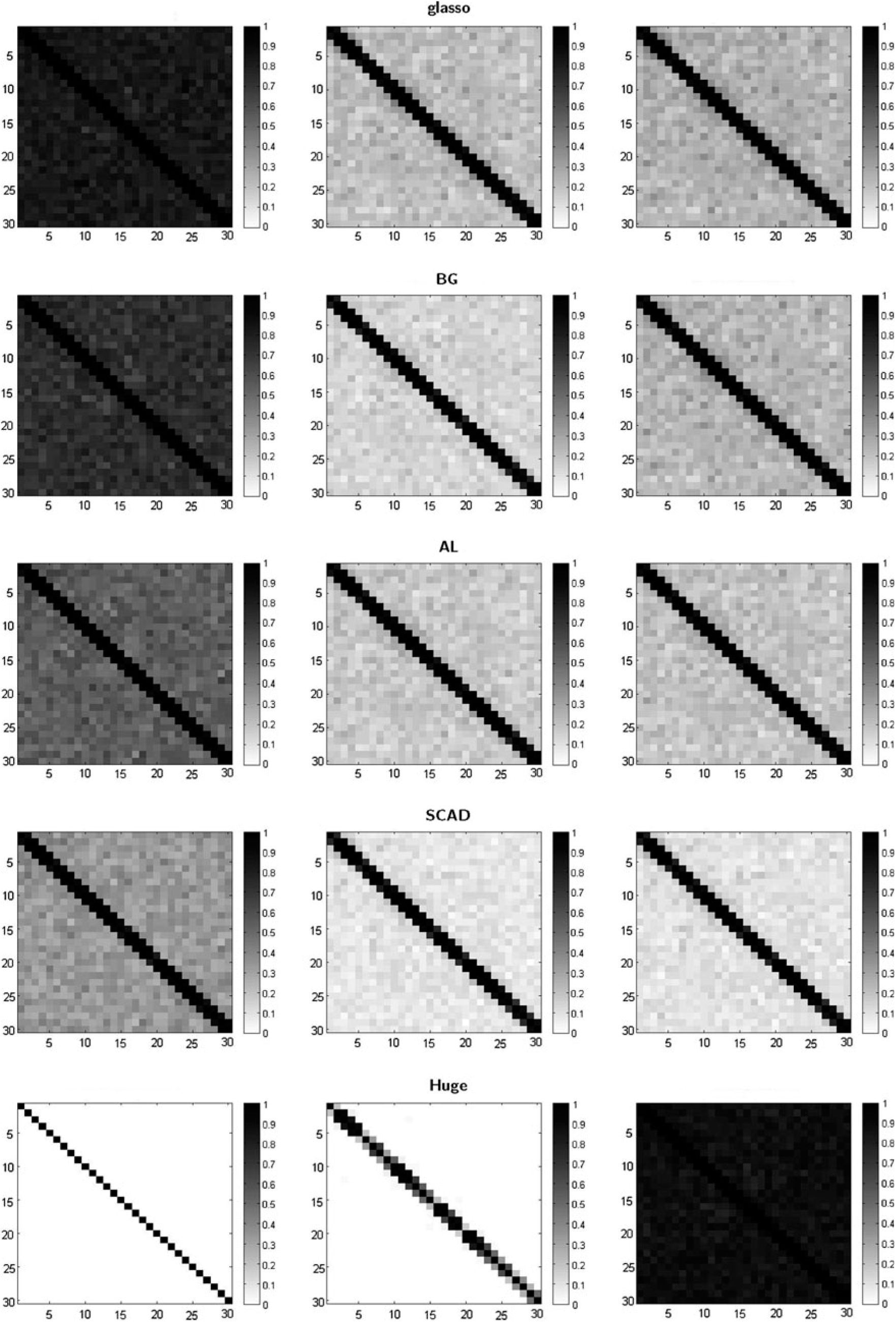
The ASP plots for the tridiagonal matrix with *a* = 1:7 for the MVN (*p* = 30) data. The left, middle, and right columns represent the selection criteria AIC, BIC, and CV for estimating methods glasso, BG, AL, and SCAD, and ebic, ric, and stars for Huge, respectively. SCAD, smoothly clipped absolute deviation.

DP-glasso performs very similarly to glasso, as does DPBG to BG, DP-AL to AL, and DP-SCAD to SCAD, hence offering no improvement. Huge estimates are either excessively dense or excessively sparse. Huge*ebic shrinks the original entries of the precision matrix to such a degree that all of their estimates are zeros, and thus, it completely loses the true model structure. Since 0.425 is the largest nonzero entry of the tridiagonal matrix, the Huge*ebic ASPs indicate that an entry value of 0.425 may be not large enough to be successfully detected, and hence, Huge*ebic may not be adequate for estimating brain networks given the magnitude of partial correlations in neuroimaging. Contrarily, Huge*stars produces too many nonzero estimates; it is only able to estimate 11:6*%* of the true zero entries, which results in extremely dense estimated matrices. Huge*ric is more capable of detecting nonzero entries than Huge*ebic; it correctly estimates 58:2*%* of nonzeros. Huge*ric also has the best TN among all method combinations. As can be seen from its ASP plot, the dark areas in the first row above and below the main diagonal have relatively large nonzero values (> 0:25). This reflects the fact that the Huge*ric combination has the most potential for correctly distinguishing between zeros and nonzero entries among all three criteria for Huge. However, its capacity is restricted to relatively large nonzero entries (e.g., > 0.25) with smaller nonzero entries (e.g., less than 0.25) often being missed. Specifically, we find that if an entry is larger than 0.31, it is very likely that this entry will not be missed by Huge *ric.

Figure 4 shows the ASP plots and Table 5 the exact TP and TN for the (moderate dimensional) exponential decay matrix. Surprisingly, Huge*ebic is the best combination that balances having both a large TP and TN. It also has the closest estimated structure to the true structure but with denser off-diagonal entries. The next best combinations are SCAD *BIC and SCAD *CV: they provide very sparse estimates (largest TN), with only entries in the first row next to the main diagonal being estimated to be nonzeros (lowest TP). However, it does not capture the true exponential decay structure along the diagonal; its estimates are closer to the tridiagonal structure. SCAD *CV appears to have marginally more white space. BG *BIC and SCAD*AIC also perform adequately; they capture most of the main diagonal structure and the off-diagonal elements have a lighter shade. AL *AIC captures the true main diagonal structure well, but the offdiagonal entries are too dense. Alternatively, AL *BIC and AL *CV lead to sparse off-diagonal entries, but the main diagonal structure is not clearly defined. The DP-glasso algorithm does not improve the glasso, BG, AL, or the SCAD methods in general. Huge*ric does not return an estimated precision matrix across all *L* = 100 repetitions. Huge*stars performs adequately; it identifies most of the main diagonal structure but suffers from severe overselection, resulting in extremely dense estimates.

**TABLE 5.** THE TRUE POSITIVE AND TRUE NEGATIVE FOR THE EXPONENTIAL DECAY MATRIX FOR THE MULTIVARIATE NORMAL (*p* = 30) DATA

| $\Omega$ | *Exp* | | | | | |
| --- | --- | --- | --- | --- | --- | --- |
|  | *TP* |  |  | *TN* |  |  |
|  | *AIC* | *BIC* | *CV* | *AIC* | *BIC* | *CV* |
| CM | 1 |  |  | 0 |  |  |
| PCM | 1 |  |  | 0 |  |  |
| Glasso | 0.94 | 0.671 | 0.621 | 0.111 | 0.487 | 0.549 |
| AL | 0.773 | 0.515 | 0.471 | 0.393 | 0.727 | 0.773 |
| SCAD | 0.677 | 0.46 | 0.415 | 0.627 | 0.887 | 0.926 |
| DP |  |  |  |  |  |  |
| Glasso | 0.94 | 0.669 | 0.621 | 0.11 | 0.488 | 0.548 |
| AL | 0.774 | 0.518 | 0.471 | 0.393 | 0.722 | 0.772 |
| SCAD | 0.677 | 0.461 | 0.417 | 0.626 | 0.889 | 0.925 |
| BG |  |  |  |  |  |  |
| *H* = 50 | 0.72 | 0.497 | 0.523 | 0.472 | 0.78 | 0.713 |
| DP-BG | 0.721 | 0.496 | 0.524 | 0.472 | 0.782 | 0.712 |
| Huge |  |  |  |  |  |  |
| Ric | NA |  |  | NA |  |  |
| Stars | 0.641 |  |  | 0.497 |  |  |
| Ebic | 0.621 |  |  | 0.781 |  |  |

**FIG. 4.**
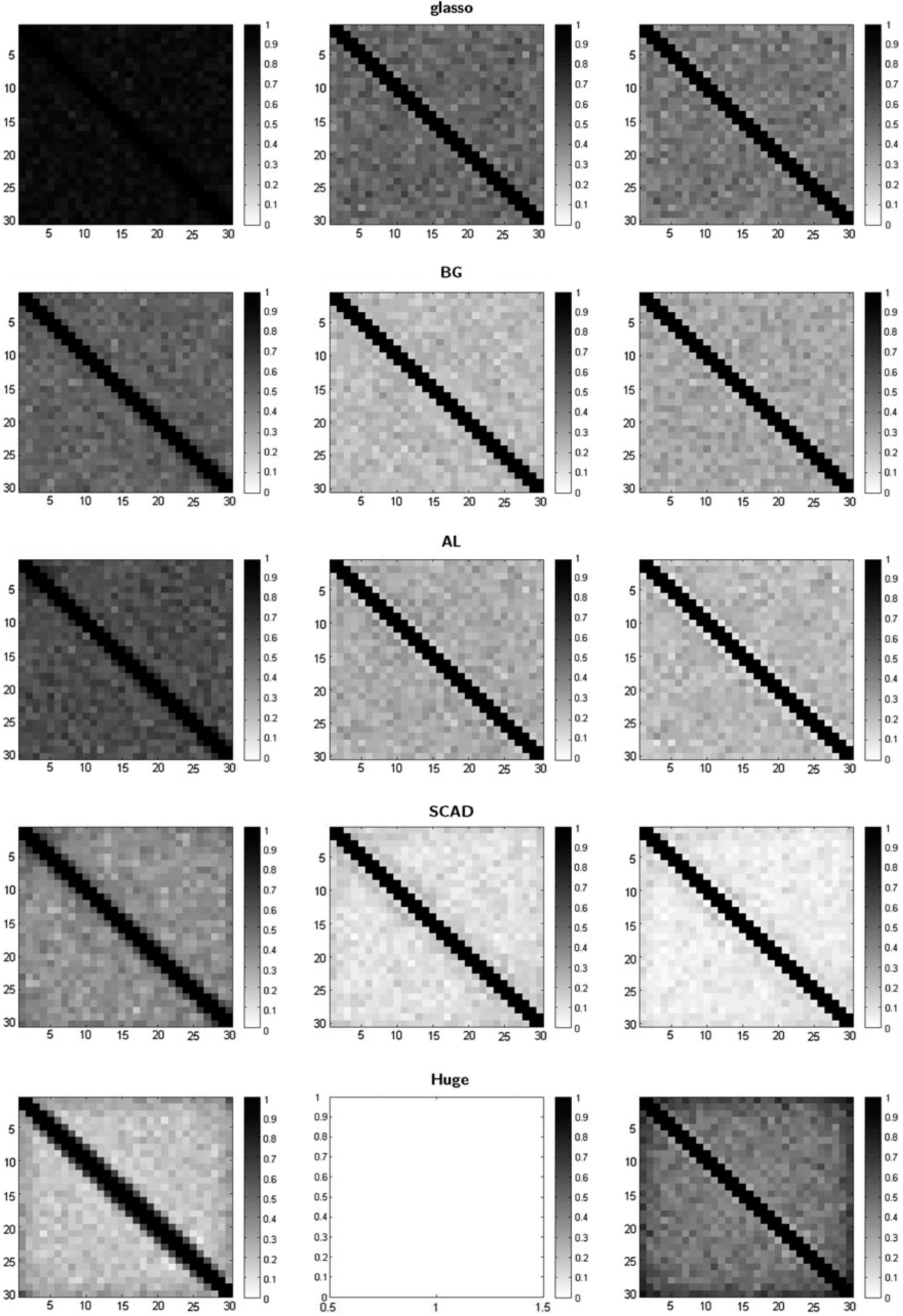
The ASP plots for the exponential decay matrix for the MVN (*p* = 30) data. The left, middle, and right columns represent the selection criteria AIC, BIC, and CV for estimating methods glasso, BG, AL, and SCAD, and ebic, ric, and stars for Huge, respectively.

Figure 5 corresponds to the ASP plots, and Table 6 the exact TP and TN for the MVN data with *p* = 30 generated from the general precision matrix. Most of the nonzero entries of this general matrix are less than 0.1 with its largest value being 0:621, and all entries are randomly spread. The best combinations are SCAD*BIC, SCAD*CV, AL*CV, AL*BIC, BG*BIC, BG*CV, and Huge*ebic but for very different reasons. SCAD*BIC and SCAD*CV provide the sparsest estimates, with around 24*%* detections of the nonzeros and 88*%* for the zero entries, providing nonzero estimates only for the relatively large entries. AL*CV, AL*BIC, and BG*CV behave similarly to SCAD*BIC and SCAD*CV but have marginally more detections of the nonzeros and marginally less detections of the zero entries. BG*BIC and Huge*ebic also perform similarly but estimate less zeros but they balance the detection of nonzeros and zeros. AIC results in dense graphs, especially glasso*AIC. This is also the case for Huge*stars. Huge*ric suffers from severe underselection again with only ∼2:4*%* of the nonzeros correctly estimated. Nevertheless, almost all zero entries are detected. The DP-glasso algorithm has equivalent TP and TN as glasso, with very similar TP and TN also for DP-BG, DP-AL, and DP-SCAD.

**TABLE 6.** THE TRUE POSITIVE AND TRUE NEGATIVE FOR THE GENERAL MATRIX FOR THE MULTIVARIATE NORMAL (*p* = 30) DATA

| $\Omega$ | *Gen* | | | | | |
| --- | --- | --- | --- | --- | --- | --- |
|  | *TP* |  |  | *TN* |  |  |
|  | *AIC* | *BIC* | *CV* | *AIC* | *BIC* | *CV* |
| CM | 1 |  |  | 0 |  |  |
| PCM | 1 |  |  | 0 |  |  |
| Glasso | 0.874 | 0.443 | 0.426 | 0.15 | 0.661 | 0.683 |
| AL | 0.627 | 0.342 | 0.302 | 0.437 | 0.775 | 0.82 |
| SCAD | 0.435 | 0.248 | 0.233 | 0.649 | 0.876 | 0.893 |
| DP |  |  |  |  |  |  |
| Glasso | 0.875 | 0.443 | 0.426 | 0.149 | 0.662 | 0.683 |
| AL | 0.627 | 0.341 | 0.302 | 0.436 | 0.775 | 0.82 |
| SCAD | 0.436 | 0.249 | 0.233 | 0.645 | 0.875 | 0.893 |
| BG |  |  |  |  |  |  |
| $H=50$ | 0.566 | 0.235 | 0.325 | 0.494 | 0.87 | 0.782 |
| DP-BG | 0.566 | 0.235 | 0.325 | 0.494 | 0.87 | 0.782 |
| Huge |  |  |  |  |  |  |
| Ric | 0.024 |  |  | 0.997 |  |  |
| Stars | 0.807 |  |  | 0.22 |  |  |
| Ebic | 0.274 |  |  | 0.853 |  |  |

**FIG. 5.**
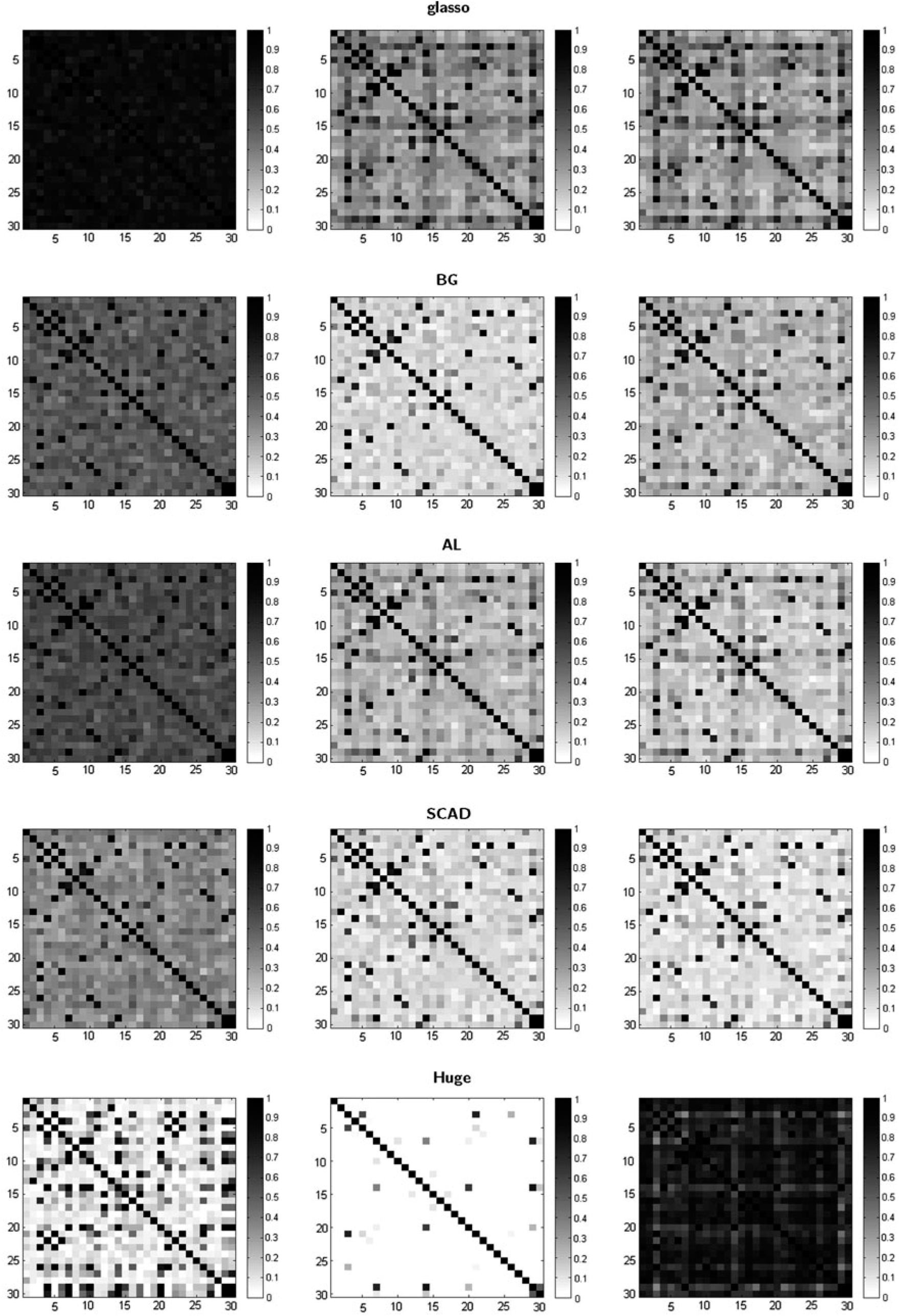
The ASP plots for the general matrix for the MVN (*p* = 30) data. The left, middle, and right columns represent the selection criteria AIC, BIC, and CV for estimating methods glasso, BG, AL, and SCAD, and ebic, ric, and stars for Huge, respectively.

### MVNAR1 data

We now apply the estimation methods in combination with the selection criteria to the MVN data with autocorrelation. To construct the autocorrelation structure, we add an AR1 model to the marginal time series from each brain region (or voxel). The results from this section should be compared to the results from the previous section (MVN data). If we fail to prewhiten the time series from each region, we obtain results similar to this section. However, if we prewhiten our time series, we obtain results similar to the previous section and thus a superior performance.

#### Low-dimensional cases

Table 7 contains the TP for the low-dimensional MVN with an AR1 autocorrelation structure added to each of its time series (MVNAR1 for short hereafter), for the four different sample sizes. As expected, compared to the MVN data, there are indeed decreases in the TP for some of the combinations. The decreases mainly occur to the AL combinations at *T* = 100, 200, SCAD combinations (especially using CV), and Huge*ric at all four different sample sizes. However, as *T* increases, the decreases in the TP become smaller, or equivalently, increasing *T* also increases the power to detect the nonzeros for autocorrelated data.

**TABLE 7.** THE TRUE POSITIVE FOR THE MULTIVARIATE NORMAL DATA WITH AR1 STRUCTURE (*p* = 5) DATA FOR FOUR DIFFERENT SAMPLE SIZES

| $\Omega$ | T = 100 | | | T = 200 | | | T = 500 | | | T = 1000 | | |
| --- | --- | --- | --- | --- | --- | --- | --- | --- | --- | --- | --- | --- |
|  | *TP* |  |  | *TP* |  |  | *TP* |  |  | *TP* |  |  |
|  | *AIC* | *BIC* | *CV* | *AIC* | *BIC* | *CV* | *AIC* | *BIC* | *CV* | *AIC* | *BIC* | *CV* |
| CM | 1 |  |  | 1 |  |  | 1 |  |  | 1 |  |  |
| PCM | 1 |  |  | 1 |  |  | 1 |  |  | 1 |  |  |
| Glasso | 1 | 1 | 0.99 | 1 | 1 | 1 | 1 | 1 | 1 | 1 | 1 | 1 |
| AL | 0.99 | 0.988 | 0.973 | 0.998 | 0.998 | 0.998 | 1 | 1 | 1 | 1 | 1 | 1 |
| SCAD | 0.9825 | 0.963 | 0.865 | 0.998 | 0.995 | 0.94 | 1 | 0.998 | 0.998 | 1 | 0.998 | 0.996 |
| BG |  |  |  |  |  |  |  |  |  |  |  |  |
| $H=50$ | 0.9875 | 0.97 | 0.978 | 0.998 | 0.998 | 0.998 | 1 | 1 | 1 | 1 | 1 | 1 |
| Huge |  |  |  |  |  |  |  |  |  |  |  |  |
| Ric | 0.393 |  |  | 0.419 |  |  | 0.648 |  |  | 0.938 |  |  |
| Stars | 1 |  |  | 1 |  |  | 1 |  |  | 1 |  |  |
| Ebic | 1 |  |  | 1 |  |  | 1 |  |  | 1 |  |  |

Table 8 contains the TN for the low-dimensional MVNAR1 data for the four sample sizes. When these results are compared with the results for the MVN data for all sample sizes (Table 3), it is evident that all of the estimating methods in combination with selection criteria AIC and BIC have smaller TN. For a number of combinations, the TN is half the rate of the MVN data. These results indicate the inferiority of the estimating methods when autocorrelation is added to the data (or the data have not been prewhitened). In general, the TN for the estimating methods in combination with AIC or BIC applied to the MVNAR1 data increases as the number of time points increases. Overall, the SCAD estimating method appears to perform the best under the assumption of autocorrelation in the data, or it can be considered the most robust method to the addition of autocorrelation. Although AL provides good estimates for the MVN data with a relatively high TN, the results for AL applied to the autocorrelated data have the largest decline in TN, especially for the AIC selection criterion. However, AL’s TN is still superior to glasso’s TN. Moreover, while both AL and BG provide similar results in terms of TN for the MVN data when *T* is large, BG provides far superior results than AL for the MVNAR1 data when they are combined with AIC and BIC. This may be due to the fact that BG bootstraps the data several times, thus removing the autocorrelation structure. Therefore, for the lowdimensional case, SCAD has the best performance, followed by BG, AL, and glasso.

**TABLE 8.** THE TRUE NEGATIVE FOR THE MULTIVARIATE NORMAL DATA WITH AR1 STRUCTURE (*p* = 5) DATA FOR FOUR DIFFERENT SAMPLE SIZES

| $\Omega$ | T = 100 | | | T = 200 | | | T = 500 | | | T = 1000 | | |
| --- | --- | --- | --- | --- | --- | --- | --- | --- | --- | --- | --- | --- |
|  | TN |  |  | TN |  |  | TN |  |  | TN |  |  |
|  | AIC | BIC | CV | AIC | BIC | CV | AIC | BIC | CV | AIC | BIC | CV |
| CM | 0 |  |  | 0 |  |  | 0 |  |  | 0 |  |  |
| PCM | 0 |  |  | 0 |  |  | 0 |  |  | 0 |  |  |
| Glasso | 0.083 | 0.14 | 0.387 | 0.067 | 0.125 | 0.367 | 0.083 | 0.17 | 0.313 | 0.074 | 0.173 | 0.263 |
| AL | 0.192 | 0.288 | 0.675 | 0.202 | 0.333 | 0.707 | 0.247 | 0.388 | 0.677 | 0.275 | 0.427 | 0.683 |
| SCAD | 0.358 | 0.518 | 0.858 | 0.386 | 0.593 | 0.902 | 0.532 | 0.668 | 0.846 | 0.777 | 0.845 | 0.885 |
| BG |  |  |  |  |  |  |  |  |  |  |  |  |
| H=50 | 0.314 | 0.473 | 0.528 | 0.335 | 0.518 | 0.501 | 0.369 | 0.544 | 0.418 | 0.339 | 0.556 | 0.382 |
| Huge |  |  |  |  |  |  |  |  |  |  |  |  |
| Ric | 0.905 |  |  | 0.882 |  |  | 0.830 |  |  | 0.568 |  |  |
| Stars | 0.145 |  |  | 0.17 |  |  | 0.238 |  |  | 0.288 |  |  |
| Ebic | 0.204 |  |  | 0.228 |  |  | 0.243 |  |  | 0.288 |  |  |

In terms of selection criteria for the MVNAR1 data, BIC correctly estimates more true zeros than AIC, however, CV outperforms both AIC and BIC. Indeed, glasso*CV at all sample sizes, AL*CV at *T* = 200, and SCAD*CV at *T* = 100, 200 have higher TN for the MVNAR1 data than for the MVN data. This indicates that if the data contain autocorrelation, which violates the independent assumption for all the estimating methods, CV combinations detect more zero entries in the precision matrices than the MVN data, for some estimating methods. By cross-validating the data, the autocorrelation structure is diluted, thus in some cases providing results very similar to the best combinations for the MVN data. However, interestingly, the BG*CV combination has lower TN at all number of time points for the MVNAR1 data compared with the MVN data, indicating that the resampling procedure neutralizes the advantage of CV.

Oddly, the TN for Huge*ric is smaller for the MVN data compared with the MVNAR1 data and Huge*ric TN decreases as the sample size increases. However, Huge*ric’s performance competes with the other best combinations for the MVNAR1 data set with very high TP and TN. For Huge*ebic and Huge*stars, the TN is larger for MVN data compared with the MVNAR1 data across all sample sizes, while their TN increases as the sample size increases as expected. Huge*ric correctly estimates significantly more zero entries than Huge*ebic and Huge*stars at all sample sizes.

#### Moderate-dimensional case

Figures 6, 7, and 8 are the ASP plots and Tables 9, 10, and 11 contain the detailed TPs and TNs for the moderate-dimensional MVNAR1 data with a tridiagonal, exponential decay and general structure, respectively. The tables show identical conclusions to their corresponding ASP plots. In this case, all the estimating methods in combination with the selection criteria AIC and BIC become significantly denser and lose the structure compared to the MVN data. While BIC can provide both accurate and sparse estimates for the moderate-dimensional MVN data, when the time series contain autocorrelation the results deteriorate dramatically, with the estimated matrices almost as dense as the matrix estimated using the AIC selection criterion. It is also evident from the ASP plots that even BIC loses some of the true graphical structure, even for the tridiagonal matrix, which contains many large entries in the precision matrix. For all the AIC/BIC combinations, SCAD is the only estimating method that does not completely lose the actual graphical model structure. However, only the basic graphical structure can be captured by the SCAD method. For all the estimating methods, the TP for the MVNAR1 data is comparable with the MVN data, but the TN declines significantly.

**TABLE 9.** THE TRUE POSITIVE AND TRUE NEGATIVE FOR THE TRIDIAGONAL MATRIX WITH *a* = 1:7 FOR THE MULTIVARIATE NORMAL DATA WITH AN AR1 STRUCTURE (*p* = 30)

| $\Omega$ | *Tri a = 1.7* | | | | | |
| --- | --- | --- | --- | --- | --- | --- |
|  | *TP* |  |  | *TN* |  |  |
|  | *AIC* | *BIC* | *CV* | *AIC* | *BIC* | *CV* |
| CM | 1 |  |  | 0 |  |  |
| PCM | 1 |  |  | 0 |  |  |
| Glasso | 0.991 | 0.982 | 0.62 | 0.049 | 0.078 | 0.792 |
| AL | 0.951 | 0.944 | 0.551 | 0.217 | 0.253 | 0.857 |
| SCAD | 0.87 | 0.844 | 0.397 | 0.483 | 0.545 | 0.945 |
| BG |  |  |  |  |  |  |
| $H = 50$ | 0.943 | 0.936 | 0.6 | 0.247 | 0.269 | 0.849 |
| Huge |  |  |  |  |  |  |
| Ric | NA |  |  | NA |  |  |
| Stars | 0.976 |  |  | 0.123 |  |  |
| Ebic | 0 |  |  | 1 |  |  |

**TABLE 10.**
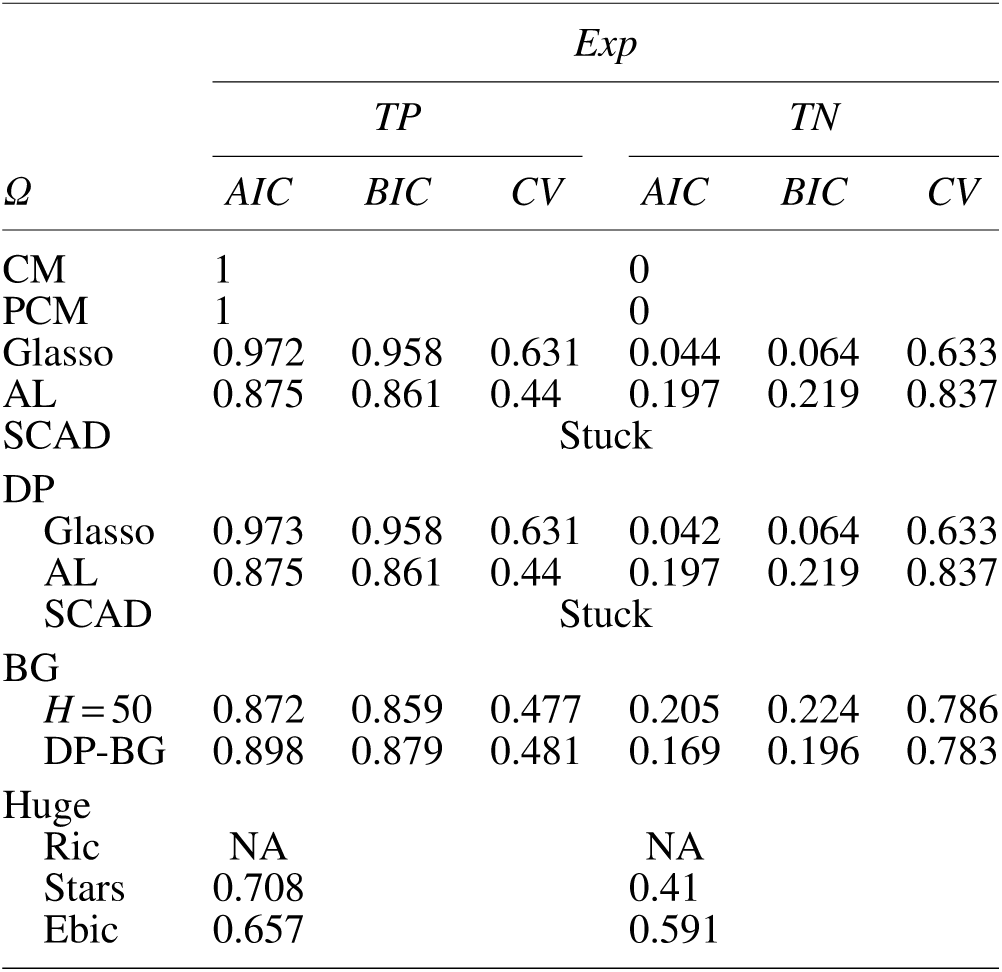
THE TRUE POSITIVE AND TRUE NEGATIVE FOR THE EXPONENTIAL DECAY MATRIX FOR THE MULTIVARIATE NORMAL DATA WITH AN AR1 STRUCTURE (*p* = 30)

| $\Omega$ | *Exp* | | | | | |
| --- | --- | --- | --- | --- | --- | --- |
|  | *TP* |  |  | *TN* |  |  |
|  | *AIC* | *BIC* | *CV* | *AIC* | *BIC* | *CV* |
| CM | 1 |  |  | 0 |  |  |
| PCM | 1 |  |  | 0 |  |  |
| Glasso | 0.972 | 0.958 | 0.631 | 0.044 | 0.064 | 0.633 |
| AL | 0.875 | 0.861 | 0.44 | 0.197 | 0.219 | 0.837 |
| SCAD |  |  |  | Stuck |  |  |
| DP |  |  |  |  |  |  |
| Glasso | 0.973 | 0.958 | 0.631 | 0.042 | 0.064 | 0.633 |
| AL | 0.875 | 0.861 | 0.44 | 0.197 | 0.219 | 0.837 |
| SCAD |  |  |  | Stuck |  |  |
| BG |  |  |  |  |  |  |
| $H = 50$ | 0.872 | 0.859 | 0.477 | 0.205 | 0.224 | 0.786 |
| DP-BG | 0.898 | 0.879 | 0.481 | 0.169 | 0.196 | 0.783 |
| Huge |  |  |  |  |  |  |
| Ric | NA |  |  | NA |  |  |
| Stars | 0.708 |  |  | 0.41 |  |  |
| Ebic | 0.657 |  |  | 0.591 |  |  |

**TABLE 11.**
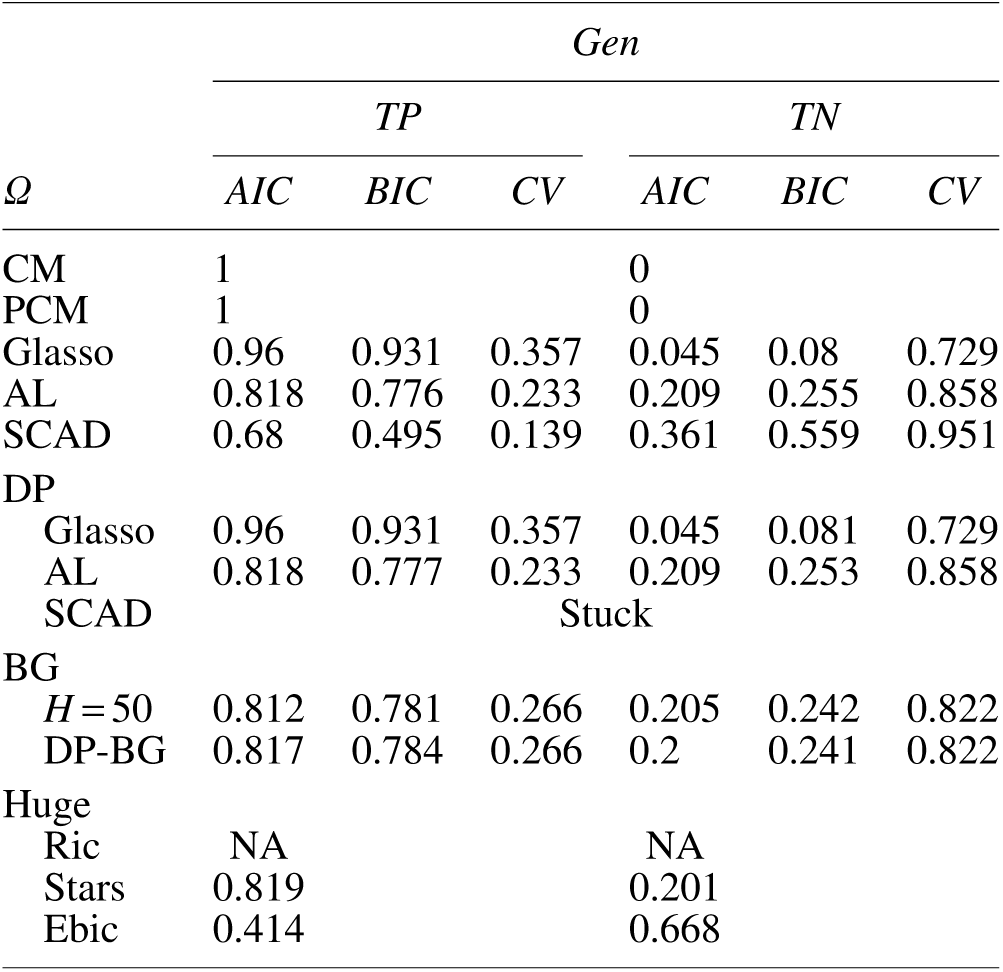
THE TRUE POSITIVE AND TRUE EGATIVE FOR THE GENERAL MATRIX FOR THE MULTIVARIATE NORMAL DATA WITH AN AR1 STRUCTURE (*p* = 30)

**FIG. 6.**
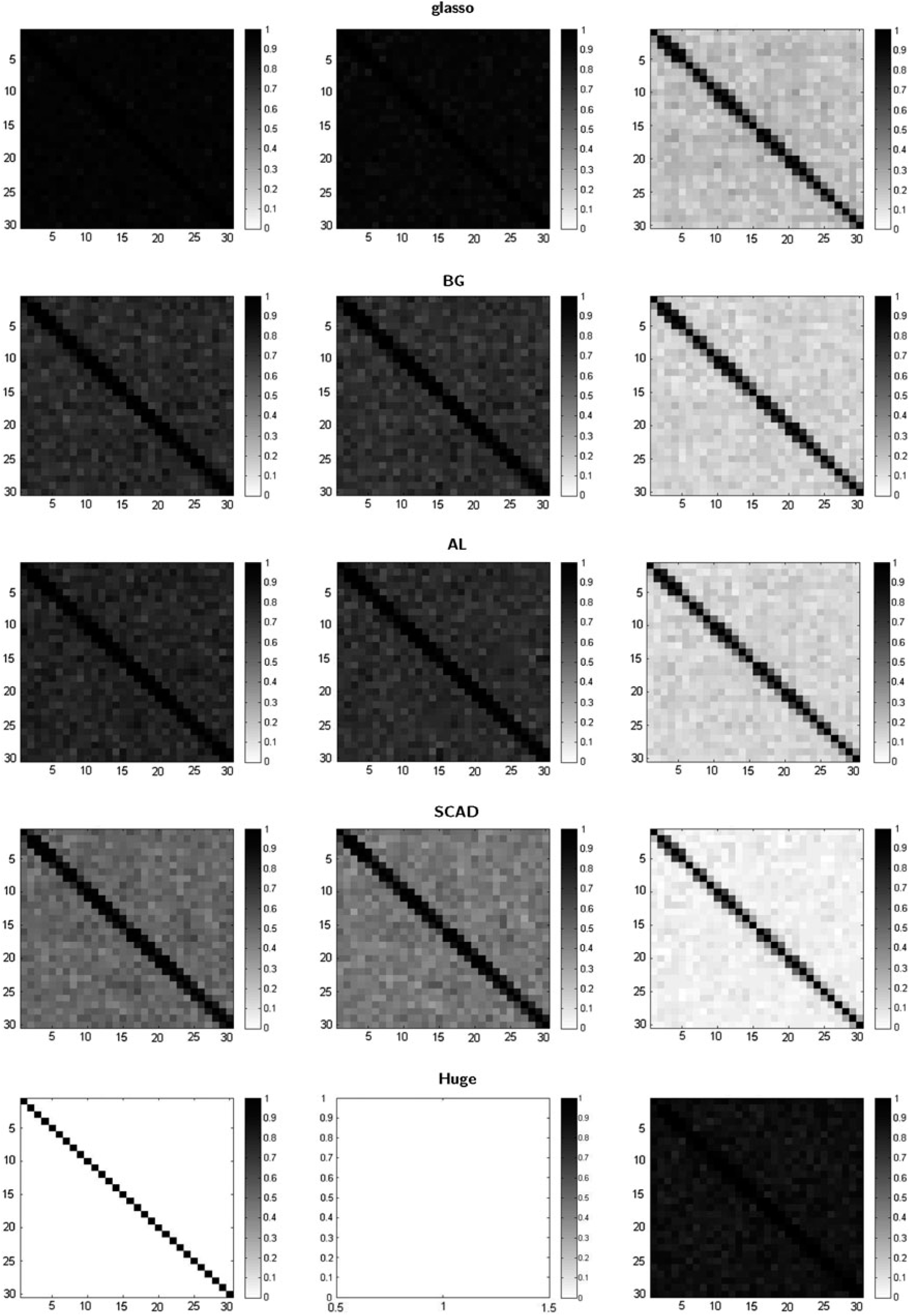
The ASP plots for the tridiagonal matrix with *a* = 1:7 for the MVN data with AR1 structure (*p* = 30) data. The left, middle, and right columns represent the selection criteria AIC, BIC, and CV for estimating methods glasso, BG, AL, and SCAD, and ebic, ric, and stars for Huge, respectively.

**FIG. 7.**
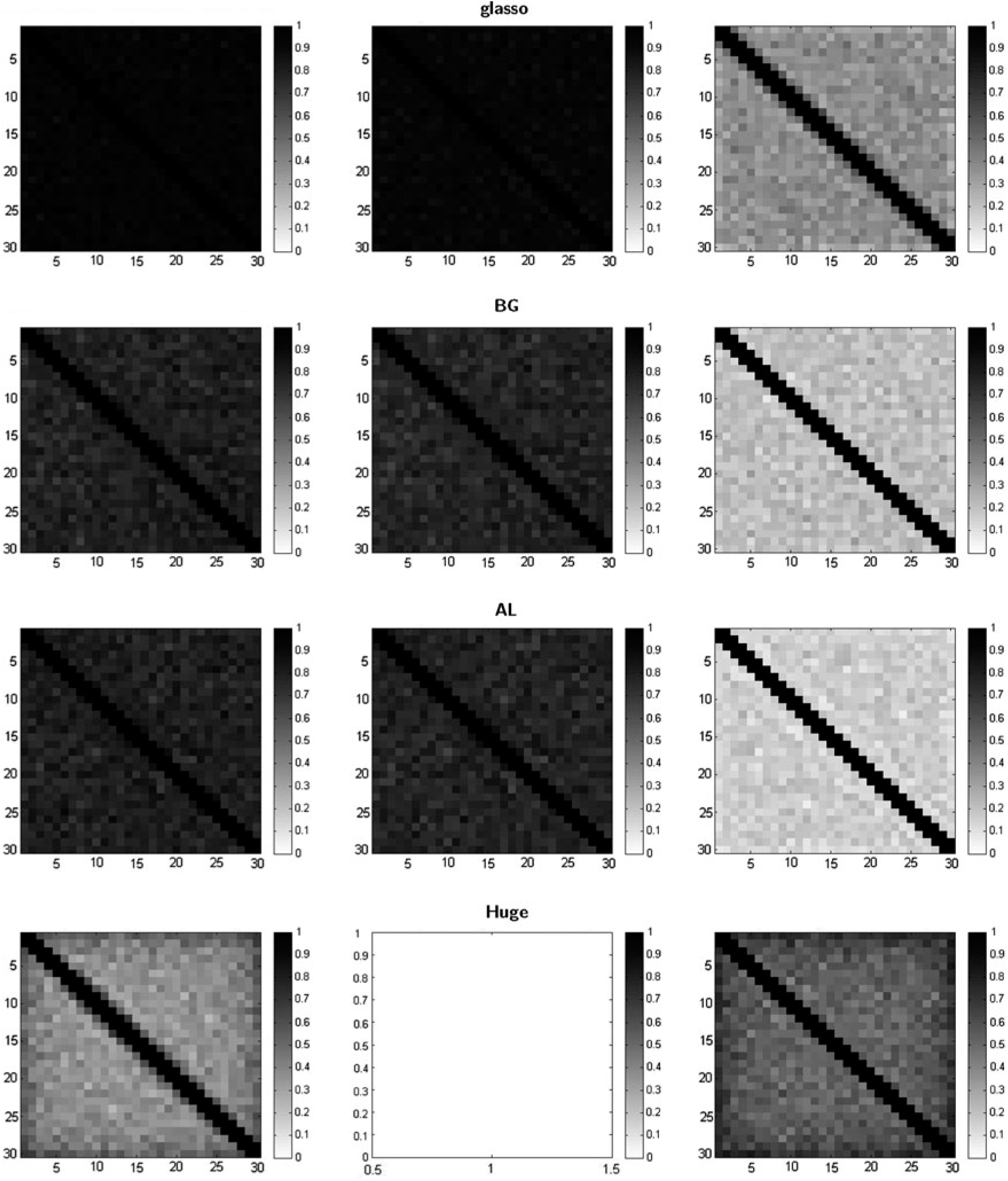
The ASP plots for the exponential decay matrix for the MVN data with AR1 structure (*p* = 30) data. The left, middle, and right columns represent the selection criteria AIC, BIC, and CV for estimating methods glasso, BG, and AL and ebic, ric, and stars for Huge, respectively.

**FIG. 8.**
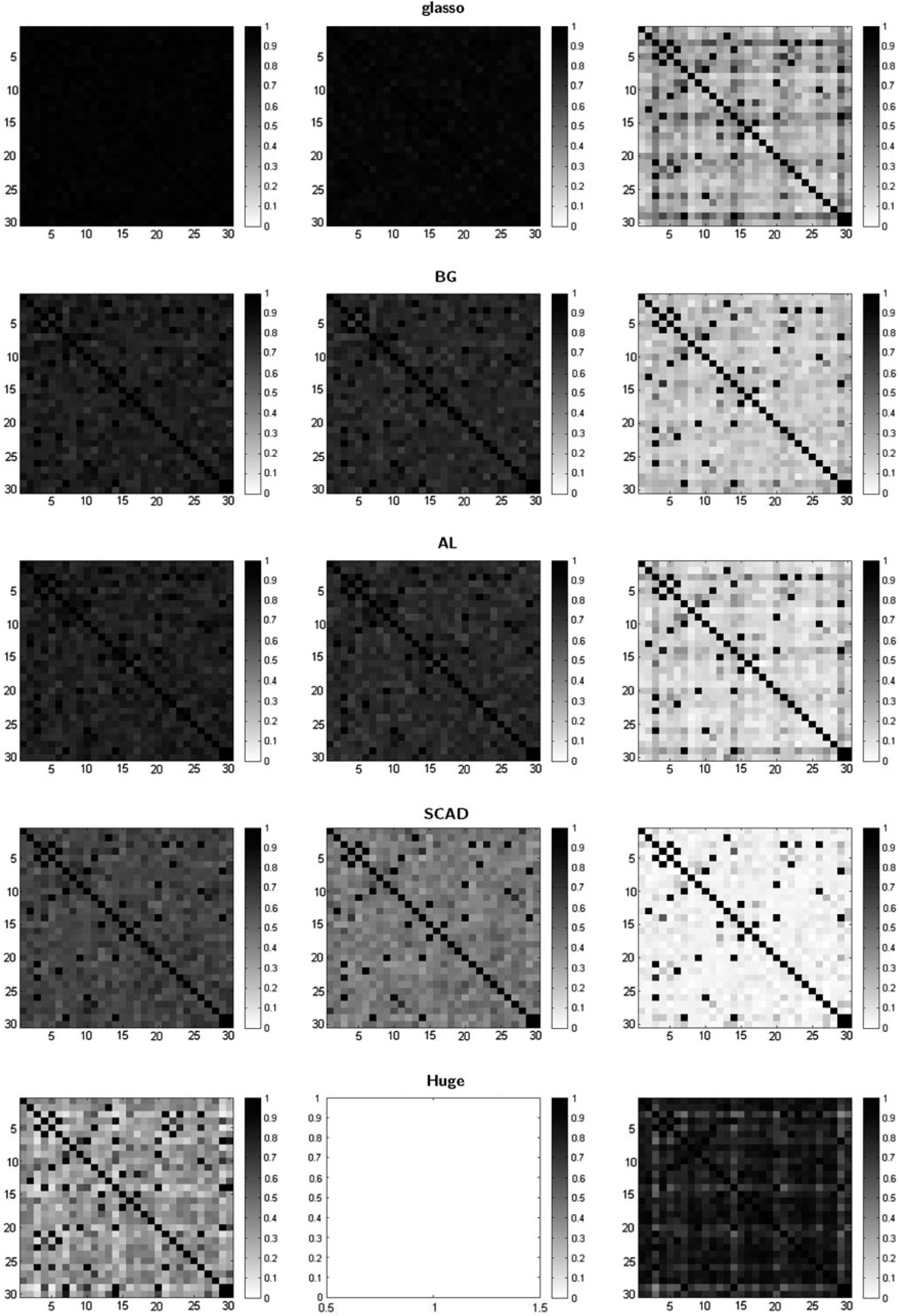
The ASP plots for the general matrix for the MVN data with AR1 structure (*p* = 30) data. The left, middle, and right columns represent the selection criteria AIC, BIC, and CV for estimating methods glasso, BG, AL, and SCAD, and ebic, ric, and stars for Huge, respectively.

Interestingly, we obtain the opposite effect for the CV selection criterion; almost all of the TPs decrease and all of the TNs increase for all the three precision matrices, leading to sparser estimates for the MVNAR1 data when compared with the MVN data. Surprisingly, this pattern even occurs for BG. For the tridiagonal matrix with *a* = 1:7, the ability of CV to detect nonzeros declines significantly. It not only declines more than both the AIC and BIC, but it also declines the most among all three precision matrix examples, which indicates that when the data are not independent, CV is more inclined to overshrink the entries of the precision matrix, even for relatively large entries. This confirms again that CV estimates are more likely to result in sparser estimates of the graph. Moreover, across all CV estimates, the SCAD*CV combination is still the sparsest.

The decline in performance of BG*CV applied to MVNAR1 data compared with MVN data in the low-dimensional cases is more significant for large sample sizes than small sample sizes. For the moderate-dimensional cases, BG*CV for the MVNAR1 data set outperforms the MVN data set. Thus, with sufficiently large sample sizes, the advantages of BG*CV cannot remedy the disadvantage of violating the independent data assumption. In other words, when the ratio of sample size to dimension is large enough, the advantage of having normal data is superior to the dilution effect of BG*CV. However, when the ratio is moderate, BG*CV is an effective way to remedy the violation of the independent data assumption.

For the Huge estimation method on the MVNAR1 data, the Huge*ebic combination performs similarly when applied to the MVN data, but the estimates remain excessively dense for the exponential decay and general matrices but extremely sparse for the tridiagonal matrix. No estimated precision matrices from the package Huge in R were returned from Huge*ric for all the 100 repetitions. This may indicate that adding an autocorrelation structure to the data may disrupt the algorithm for Huge*ric. Finally, Huge*stars applied to MVNAR1 data performs similarly to the application to the MVN data, it returns very dense estimates.

### fMRI results

In Figures 9 and 10, we show that the ASP plots for the resting-state fMRI data averaged over the 45 subjects without and with prewhitening of the ROI time series, respectively. Elements 1–8, 9–10, 11–13, 14–20, 21–29, and 30–31 on the *x*-axis (and *y*-axis) of the ASP plot coincide with the 8, 2, 3, 7, 9, and 2 regions from the attentional network, the visual network, the sensorimotor network, the salience network, the default mode network, and the auditory network, respectively. Remarkably, many of the combinations capture the block diagonal structure of the networks, indicating that most of the connectivity is intra-network with some internetwork connectivity as well. For example, there is high internetwork connectivity between the attentional and salience, attentional and default mode, visual and default mode, visual and sensorimotor, visual and salience, and auditory and default mode networks. The only combinations that fail to capture this structure are glasso*AIC, glasso*BIC, Huge*ebic, and Huge*stars for the data without prewhitening and glassoAIC, glasso*BIC, glasso*CV, Huge*ric, and Huge*stars for the data with prewhitening.

**FIG. 9.**
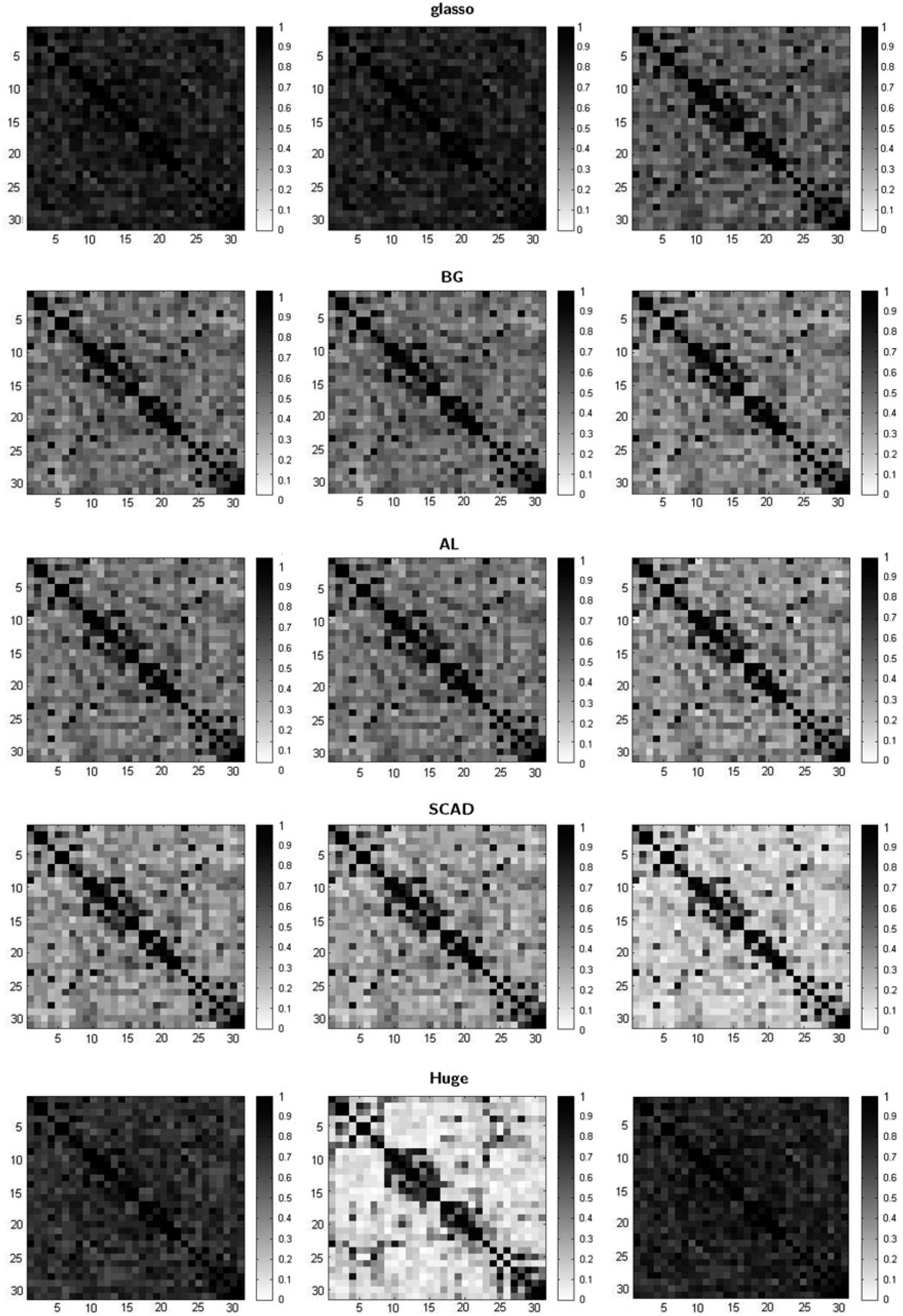
The ASP plots for the resting-state fMRI data (without prewhitening). The left, middle, and right columns represent the selection criteria AIC, BIC, and CV for estimating methods glasso, BG, and AL and ebic, ric, and stars for Huge, respectively. AIC, Akaike information criterion; AL, adaptive lasso; BG, bootstrap graphical lasso; BIC, Bayesian information criterion; CV, cross-validation; ebic, extended BIC; fMRI, functional magnetic resonance imaging; Huge, high-dimensional undirected graph estimation; ric, rotation information criterion; stars, stability approach for regularization selection.

**FIG. 10.**
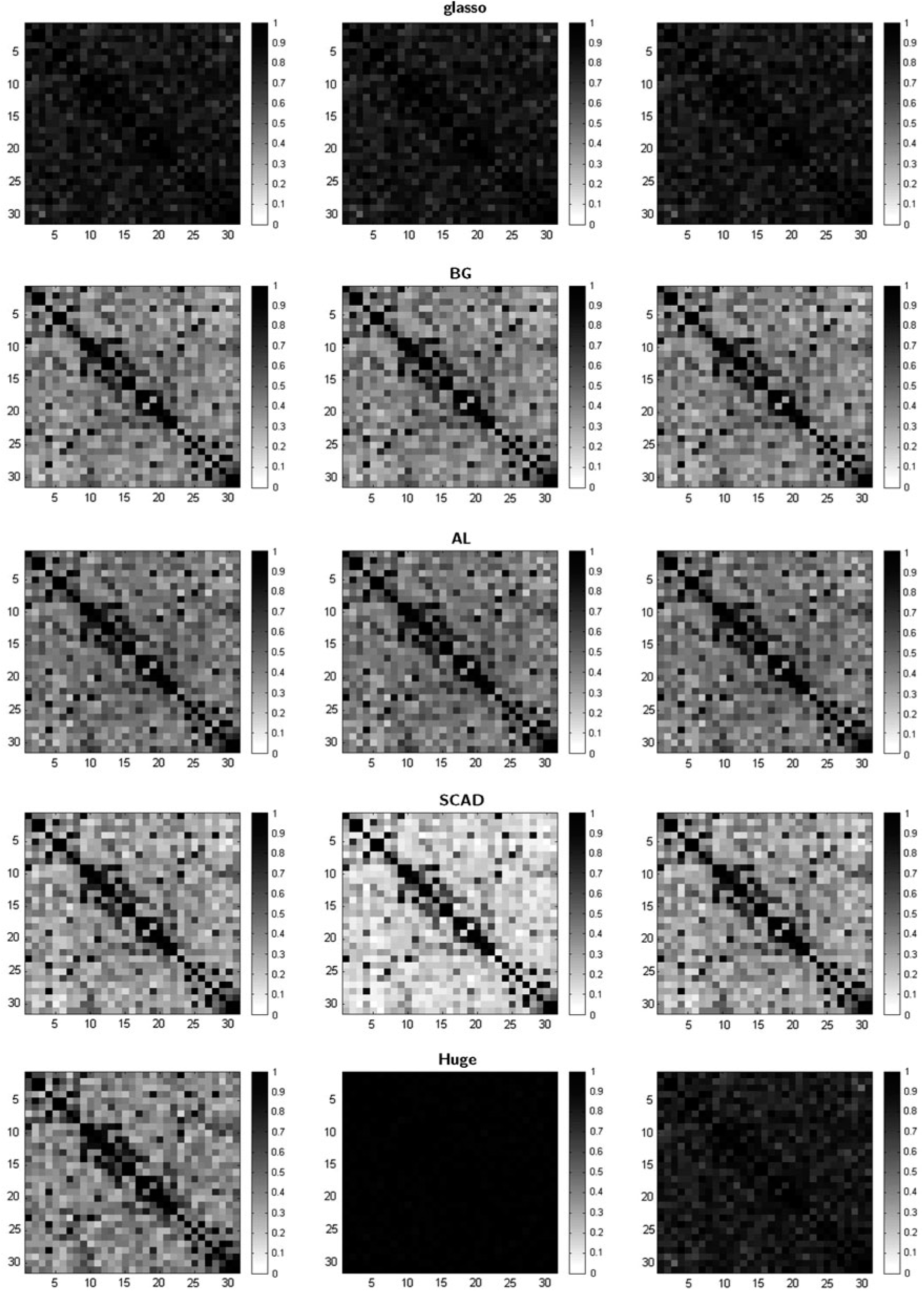
The ASP plots for the resting-state fMRI data (with prewhitening). The left, middle, and right columns represent the selection criteria AIC, BIC, and CV for estimating methods glasso, BG, and AL and ebic, ric, and stars for Huge, respectively.

The ASP plots of the prewhitened ROI time series and the ROI time series with autocorrelation are quite similar in terms of structure and density apart from the Huge and glasso combinations. This is unexpected given that in the simulation study, we found that the TP is higher for the data with autocorrelation compared with the independent data, while the TN declines significantly. For the ROI time series with autocorrelation (Fig. 9), BG, AL, and SCAD are the estimating methods that clearly capture the block diagonal structure of the network across all three selection criteria, AIC, BIC, and CV. However, the estimates of SCAD are slightly less dense than BG and AL. It is clear that the combination SCAD*CV is best overall as it depicts a clear block diagonal structure that is sparse. The reason for its superior performance is that for CV, the TP decreases and the TN increases leading to sparser estimates for the data with autocorrelation as CV dilutes the autocorrelation structure. This is consistent with our simulation study. Huge*ebic also captures the block diagonal structure, while its combination with ric and stars produces very dense structures. glasso*AIC and glasso*BIC produce excessively dense results, which almost lose the basic network structure of the data set.

For the ROI time series that have been prewhitened (Fig. 10), BG, AL, and SCAD in combination with AIC, BIC, and CV capture the block diagonal structure of the network. The SCAD*BIC combination produces the best results for this data set, producing the sparsest estimate with a clear block diagonal structure. Again, this is consistent with our simulation results. Huge*ric somewhat captures the block diagonal structure, while its combination with ebic and stars produces very dense structures. On the contrary, Huge*ric and Huge*stars and all three glasso combinations fail to detect the block diagonal structure of the data set, leading to excessively dense estimates.

## Discussion

### Computation

In the Results section, we considered the performance of the sparse network estimating methods in combination with selection criteria in terms of correctly estimated nonzero (TP) and zero (TN) elements of the precision matrices. We now discuss the computational cost of the combinations, which is also a very important practical criterion. To compare the computational time for each combination *M*_*q*_ * *C*_*c*_, we fixed the repetition time (*L* = 100), the number of time points *9T* = 100), the dimension (*p* = 5), the data type (MVN), and the path of the regularization parameters (*ρ*) for each combination to be *ρ* = 0:01 × *i* for *i* = 1, …, 100. In addition, for the BG and DP-BG estimating methods, we fixed the resampling number to be *H* = 50 and the threshold to be *π*_*thr*_ = 0:9. Table 12 shows the computational time (in sec) for each combination, *M*_*q*_ * *C*_*c*_, to complete *L* = 100 repetitions of each combination, *M*_*q*_ * *C*_*c*_. Specifically, it is the time to estimate the true precision matrix *Ω* based on *N* = 100 different data sets by estimating method *M*_*q*_ and then selecting the best estimate by selection criterion *C*_*c*_ among all of the 100 estimated precision matrices produced by the 100 regularization parameters. The glasso algorithm is by far the most computationally efficient estimating method, but combining computational cost and the performance of the estimating method, both SCAD and AL in combination with BIC perform best. While BG performs well, it is very slow computationally given that it has to resample the data many times and estimate the precision matrix using glasso. The DP algorithms are also considerably less efficient across all estimating methods.

**TABLE 12.** THE COMPUTATIONAL TIME (IN SEC) OF EACH COMBINATION APPLIED TO THE MULTIVARIATE NORMAL (*p* = 5) DATA USING AN INTEL CORE I3-2350 M 2.30 GHZ CPU

| Methods | Time | Methods | Time |
| --- | --- | --- | --- |
| Glasso*AIC | 3.44 | DP-G*AIC | 76.8 |
| Glasso*BIC | 3.44 | DP-G*BIC | 76.66 |
| Glasso*CV | 37.48 | DP-G*CV | 394.09 |
| BG*AIC | 170.31 | DP-BG*AIC | 3973.94 |
| BG*BIC | 171.83 | DP-BG*BIC | 3971.14 |
| BG*CV | 2021.85 | DP-BG*CV | 24129.8 |
| AL*AIC | 8.65 | DP-AL*AIC | 80.84 |
| AL*BIC | 9.17 | DP-AL*BIC | 80.64 |
| AL*CV | 80.35 | DP-AL*CV | 431.15 |
| SCAD*AIC | 15.82 | DP-SCAD*AIC | 82.34 |
| SCAD*BIC | 14.57 | DP-SCAD*BIC | 82.27 |
| SCAD*CV | 107.16 | DP-SCAD*CV | 432.75 |
| Huge*ric | 115.64 | Huge*ebic | 87.77 |
| Huge*stars | 1981.21 |  |  |

### BG algorithm parameter choices

Next, we consider the optimal choices for the parameters of the BG: the number of bootstrap resamples *H* and the threshold probability *π*_*thr*_. Since we obtain TP = 1 across almost all methods for the simulated low-dimensional MVN data set, we concentrate our comparison on the TN of the methods. Table 13 illustrates the TN of glasso and BG in combination with AIC, BIC, and CV, with the threshold π_*thr*_ being set to values 0:75, 0:8, 0:85, 0:9, 0:95. For each threshold value, *π*_*thr*_, we also consider four different repetition values for the number of resamples, *H* = 50, 100, 150, 200 with a fixed number of time points *T* = 200.

**TABLE 13.**
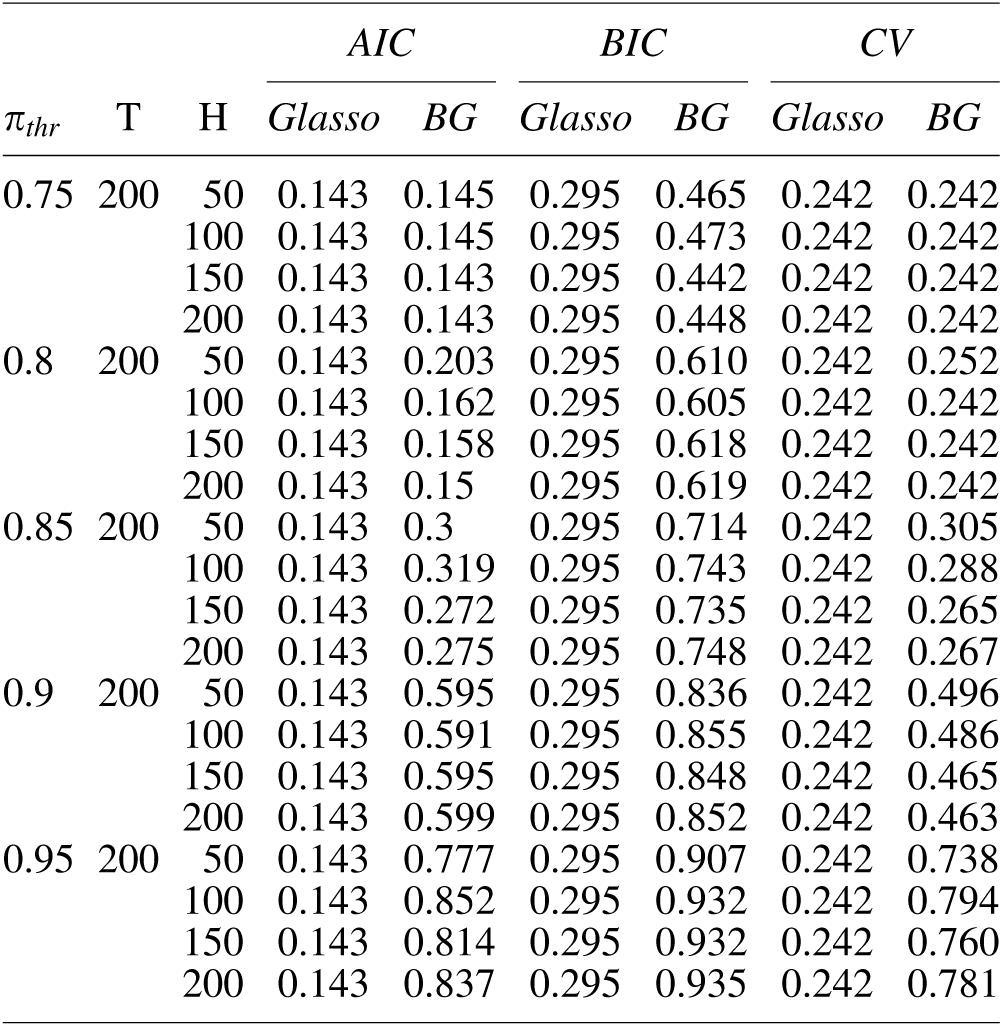
THE TRUE NEGATIVE FOR THE MULTIVARIATE NORMAL (*p* = 5) DATA AND *T* = 200 USING GLASSO AND BOOTSTRAP GLASSO FOR *H* = 50, 100, 150, 200, *π*_*thr*_ = 0:75, 0:8, 0:85, 0:9, 0:95, AND SELECTION CRITERIA AIC, BIC, AND CV

#### How *H* affects BG

From Table 13, it is evident that increasing the number of resamples from 50 to 200 provides no improvement in the estimates of BG. In some cases, having too many resamples adversely affects the results. We found that the number of resamples, *H*, should not be greater than the number of time points, *T*. Indeed, a larger *H* leads to markedly more computational time, especially for moderate-dimensional data sets. In conclusion, increasing *H* can marginally boost the precision of the estimate, but it also significantly increases the computational time.

In our simulations, we fixed the number of time points, *T* = 100, in our moderate-dimensional data sets and considered *T* = 100, 200, 500, 1000 in our low-dimensional data sets, and set *H* = 50 for BG. Hence, *H* is at most half the number of time points, but it still effectively improved the estimates over the glasso estimates, however, it required additional computational time. In addition, *H* = 50 may be not large enough for data sets with a large number of time points. Indeed, for data sets with a large number of time points, we believe that increasing *H* will produce better estimates for BG, but we do not think the trade-off in the accuracy of the estimates is offset by increasing the number of resamples and thus the computational time.

How *π*_*thr*_ affects BG. From Table 13, it is evident that *π*_*thr*_ has more influence on the BG estimates than the number of resamples, *H*. A marginal increase in the value of *π*_*thr*_ leads to a major improvement in the estimates. However, when an excessively large *π*_*thr*_ is set, the nonzero estimate frequencies struggle to become larger than the threshold, thus reducing the TP. Thus, many estimated entries are set to zero and the estimated graphs are very sparse with many false negatives. Another appealing feature of *π*_*thr*_ is that no additional computational time is required. As the best performance (balancing TP and TN) is found using *π*_*thr*_ = 0:9, we consequently choose *π*_*thr*_ = 0:9 in all our BG simulations, which effectively improves the estimates without the risk of providing inordinately sparse results.

### Accuracy comparisons

We now discuss the precision of the estimating methods for the data sets in the Simulations section, including the MVN and MVNAR1 data. We consider the accuracy and the sparsity levels of the methods, which are not equivalent to each other. For example, an estimate can be very sparse, but it may be unnecessarily sparse in that it cannot effectively reveal the true graphical structure, which is regarded as undesirable. Conversely, an estimate may be denser than another, but if it depicts the true graphical structure, it can be regarded as the better estimate. In addition, sparsity and the accuracy are sometimes equivalent because nonzero entries are easier to estimate than zero entries. Thus, provided that the majority of nonzeros are estimated correctly, a sparser graph indicates that more zero entries have been estimated correctly, which leads to a more accurate estimate that is closer to the true graphical structure.

The results in the simulation study represent the best estimated undirected graph under the conditions that certain parameters were prespecified, such as the path of regularization parameters (100 *ρ*s), the number of bootstrap resamples (*H*), and the BG threshold (*π*_*thr*_). Accordingly, an estimation method ***M***_*q*_ is superior to another method ***M***_*l*_ if the best estimate 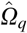 is superior to the best estimate 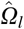. We also make the same conclusions for selection criteria. As our results and conclusions are based on the parameters we have chosen, it is possible that superior results could be found if some parameters are adjusted. However, the results may also deteriorate if some undesirable parameters are chosen, such as choosing a narrow range of *ρ*s, using only one fixed *ρ*, using a *π*_*thr*_ that is too small, or using an inferior number of resamples, *H*.

#### Computational highlights

In general, glasso is a simple and an efficient algorithm for estimating a sparse precision matrix, *Ω*. In addition, the popular R package glasso (or “glmnet” in MATLAB) makes it very convenient to implement. A very attractive feature of glasso is its desirable computational speed. For example, it has been shown that based on an Intel Xeon 2.80GH processor with 2 to 8 iterations of the outer loop, *p* = 400 in a sparse *Σ* case, it only takes glasso 1.23 sec (the CPU time spent in the C program since glasso was coded in Fortran and linked to an R language function) to estimate the precision matrix (Friedman et al., 2007a). Another advantage of glasso is that the positive definiteness of each updated *Σ* is ensured.

The BG method combines an initial estimation method, glasso, with the bootstrap, which provides many desirable properties. We found that the results for BG vary marginally with a different number of resamples, *H*, but vary considerably with different threshold parameters, *π*_*thr*_. The percentage of false estimates (estimating nonzeros for zeros or zeros for nonzeros) is controlled or bounded. BG provides improvements in accuracy over the glasso method. However, a major disadvantage is that it is more time-consuming than glasso.

The newly developed DP-glasso algorithm is similar to glasso, except that its optimization variable is the precision matrix rather than the covariance matrix. Beginning with any positive definite matrix, DP-glasso produces a sparse and positive definite precision matrix. However, for glasso and DP-glasso, one member of the pair (*Ω, Σ*) is not the inverse of the other. Moreover, Mazumder and Hastie (2012) found that, not only theoretically but also experimentally, the DP-glasso is computationally more efficient than glasso. However, our results did not have this conclusion. Overall, Huge’s performance was poor. In general, it found excessively dense or sparse networks. Perhaps this may be due to the dimension of our simulations, given that Huge was motivated using (very) high-dimensional data sets.

#### Conclusions for the MVN data

In both the lowdimensional and moderate-dimensional cases, when *T* is of moderate size, SCAD*BIC or SCAD*CV performs the best. When *T* is large, we recommend using SCAD*BIC, SCAD*CV, AL*BIC, AL*CV, and BG*BIC. We do not recommend using the glasso and Huge combinations for the low-dimensional settings and the glasso*AIC, Huge*ebic, and Huge*stars combinations for the moderate-dimensional settings. By increasing the dimension of the MVN data, we do not see a deterioration in the estimates using glasso, AL, BG, or SCAD. On the contrary, with increasing dimensions, we find remarkably better detections of the zero entries by glasso*BIC and glasso*CV, along with moderate increases by AL*BIC, AL*CV, and BG*CV. Huge does not perform as well as glasso, AL, BG, and SCAD as the dimension increases. Huge*ric is recommended only if the true precision matrix contains a sizeable number of large nonzero elements. Otherwise, all the three Huge combinations should not be adopted, due to their incapability of maintaining the true graphical structures.

If the data set is MVN, combinations such as SCAD*BIC, SCAD*CV, or BG*BIC provide accurate estimates. For example, more than 90*%* of all the entries of the true precision matrix can be successfully detected. Note that glasso, AL, BG, and SCAD have a superior ability to detect smaller nonzero entries than Huge. In other words, the estimation precision of glasso, AL, BG, and SCAD is better than Huge. Thus, if the connections within a graph are not very large, glasso, AL, BG, and SCAD are better choices than Huge, with SCAD being the strongest method.

#### Conclusions for the MVNAR1 data

In general, SCAD is the most stable and best estimating method for data with an autocorrelation structure. AL performs the worst in correctly estimating zero entries. CV is the most resistant selection criterion to the autocorrelation structure. Furthermore, all the estimating methods’ ability to correctly detect true entries in the precision matrix increases with the number of time points.

For the low-dimensional MVNAR1 data, we recommend SCAD*CV and AL*CV across all sample sizes. SCAD*BIC is also recommended for large *T*. For the low-dimensional MVNAR1 data, the Huge*ric combination is superior to the other combinations of Huge. Huge*ric’s power for detecting zero entries increases dramatically, while its ability for estimating nonzeros is greatly weakened, for small sample sizes. While all of the Huge combinations maintain the true graphical structure, the Hug*ebic and Huge*stars estimates are more dense than Huge*ric.

For the moderate-dimensional MVNAR1 data, AL*CV, BG*CV, and glasso*CV are favored for the matrices with relatively small entries, since SCAD*CV overshrinks entries, leading to excessively sparse estimates. If the true entries are relatively large, SCAD*CV is recommended as it achieves more sparsity than the other combinations. The Huge combinations have a different performance in the moderate-dimensional case compared to the low-dimensional case. Huge*stars and Huge*ebic still produce excessively dense and sparse estimates as they do for the other moderate-dimensional data sets.

In conclusion, the AR1 autocorrelation structure remarkably reduces the sparsity of the estimated graphs. In other words, for both lowand moderate-dimensional simulation settings, the results of the MVNAR1 data are much denser than the MVN data. In addition, the ability to correctly estimate both nonzeros and zeros becomes significantly weaker. The excessively dense graphs estimated by the majority of the estimating methods for the data with autocorrelation indicate that the autocorrelation structure inherent in the data causes the estimating methods to perform poorly. If there is autocorrelation in the data and you are not confident in the prewhitening procedure, our advice is to use the CV selection criterion. By cross-validating the data, the autocorrelation structure is diluted, and thus, in some cases providing results very similar to the best combinations for the MVN data.

#### Overall conclusions

In general, for all dimensions and types of data sets, our conclusions are as follows:

1. SCAD > AL > BG > glasso > Huge.
2. The DP-glasso algorithm is not effective in improving the estimates at least for the data sets we considered. In other words, DP-glasso has very similar results to glasso, and hence, DP-BG to BG, DP-AL to AL, and DP-SCAD to SCAD.
3. BIC always chooses sparser estimates than AIC. CV is sparser than AIC most of the time. All of the selection criteria are able to preserve the true graphical structure with the only difference between them being the sparsity of the estimated graphs.
4. Typically, SCAD*BIC provides the best estimates for all the simulated data sets. AL*BIC, BG*BIC, or AL*CV is the second-best choice most of the time. glasso*AIC and Huge always provide undesirable estimates.

#### Limitations

While our simulation study is extensive and considers various settings (different sample sizes, dimensions, data types, and sparsity levels), we acknowledge that we have not covered all possible settings. We consider two different data types, MVN and MVNAR1, to show the effect of autocorrelation on the methods. While we could have considered various degrees of autocorrelation inherent in the ROI time series in our simulation study, we thought by considering MVN and MVN with a large degree of autocorrelation (*ϕ*_1_ = 0.8 in the AR1 model), the results for the other degrees of autocorrelation would lie in the spectrum between the two extremes. Furthermore, we have not considered the fact that there exists voxel-level spatial correlation within an ROI. We believe that ignoring this correlation by simply taking the average voxel time courses in each ROI would also have an effect on TP and TN in the same way ignoring the temporal correlation (i.e., without prewhitening data) has effect on TP and TN. However, we believe that the simulation study provides a comprehensive review of the methods that has a clear message and we have concentrated on the issue of autocorrelation as this violates the assumption of the graphical models.

Finally, as fMRI data are distanced from the underlying neural sources by many confounding stages, a careful validation is necessary before safely interpreting the results of the network estimation methods (Smith et al., 2011). Typically, the closer a given data set is with the assumptions of the estimation methods, the more desirable the results are. Hence, the assumptions of the underlying model should be checked.

## Conclusion

In this work, we studied various procedures for estimating sparse brain networks. To find the optimal sparsity level for each network, we considered various selection criteria. We showed by using an extensive simulation study that the best estimating method was SCAD in combination with either BIC or CV. We also studied the effect of autocorrelation, inherent in neuroimaging data, on the estimating methods and selection criteria. Overall, we found that the presence of autocorrelation had a negative effect on the estimates and resulted in denser networks when compared to data that had been prewhitened. We hope that our work encourages neuroimaging researchers to think more carefully about the sparse graphical methods used to estimate their FC networks and the effect autocorrelation has on their estimated FC networks. It is critical that neuroscientists know the decisions they are making when estimating FC networks. In addition, although the main focus of this work is on estimating methods that estimate static FC where the time series data from each brain region are stationary, the methods can be easily incorporated into an algorithm for estimating dynamic FC via a sliding window or for estimating FC change points in a similar vein to Cribben et al. (2013, 2012) and Cribben and Yu (2017), which is a recent area of interest in the neuroimaging community. Future work entails how graph metrics (e.g., small-worldness and modularity) are affected by the estimating procedure used and the presence of autocorrelation.

## Acknowledgments

We acknowledge the efforts of all individuals responsible for designing and carrying out the experiment and for collecting the data. I.C. was supported by the Pearson Faculty Fellowship (Alberta School of Business) and an Alberta Health Services (AHS) grant.

